# A Multi-Organ Murine Metabolomics Atlas Reveals Molecular Dysregulations in Alzheimer’s Disease

**DOI:** 10.1101/2025.04.28.651123

**Authors:** Simone Zuffa, Celeste Allaband, Vincent Charron-Lamoureux, Andres M. Caraballo-Rodriguez, Abubaker Patan, Ipsita Mohanty, Julius Agongo, John W. Bostick, T. Jaymie Connerly, Taren Thron, Brittany Needam, Matheus de Castro Fonseca, Rodolfo Salido Benitez, Lauren Hansen, Helena Tubb, Jennifer Cao, Karel Kalecký, Teodoro Bottiglieri, Siamak MahmoudianDehkordi, Leyla Schimmel, Alexandra Kueider-Paisley, Stewart F. Graham, Dionicio Siegel, Mingxun Wang, Rob Knight, Rima Kaddurah-Daouk, Pieter C. Dorrestein, Sarkis K. Mazmanian, the Alzheimer Gut Microbiome Project Consortium

## Abstract

The etiology of Alzheimer’s Disease (AD) remains largely unclear but is likely driven by gene-environment interactions. Here, we present a multi-organ untargeted metabolomics dataset (2,271 samples) generated from five tissue types in two genetic AD mouse models under colonized or germ-free conditions, complemented by shotgun metagenomics sequencing data (666 samples). Systems-level analyses of 3xTg and 5xFAD mice reveal clusters of dysregulated molecular classes across tissues including carnitines, bile acids, B vitamins, and neurotransmitters. This signature, coupled with microbiome profiles, suggests increased oxidative stress via mitochondrial dysfunction. Molecular feature tracking via tissueMASST, a mass spectrometry search tool we developed to bridge animal model findings with human data, identifies microbially-modulated phenylacetyl-carnitine as positively associated with aging and cognitive impairment across human AD studies. With hundreds of yet-to-be-characterized metabolites, this public resource and its associated tools will aid future research in the pathophysiology of AD.

## INTRODUCTION

Alzheimer’s disease (AD) is the leading cause of dementia with more than 7 million cases in the United States^1^ and a projected economic burden reaching trillions of dollars in the coming years^2^. While the causes of AD remain incompletely understood, gene-environment interactions are likely involved and represent a frontier of research in the field.

The gut microbiome is a major environmental contributor to disease and regulates the production of metabolites for bioenergetics, neurotransmitters, vitamins, and numerous other yet-to-be-characterized molecules that may play a crucial role in metabolic homeostasis^3–6^. Importantly, the gut microbiome has been linked to neurodegenerative and neuropsychiatric diseases^7,8^. Thus microbial metabolites, along with immune^9^ and endocrine signaling pathways^10^, may impact vulnerability to and progression of AD. Notably, we previously implicated dysregulation of bile acids, which are metabolized by the gut microbiome, in cognitive decline and brain imaging changes in humans^11–14^ and others showed that antibiotics administration may reduce neuroinflammation^15,16^. Additionally, untargeted metabolomics and metagenomics analyses have revealed alterations in gut microbial community composition and function in AD^17–21^. Understanding how the gut microbiome contributes to AD pathophysiology may uncover aspects of the multifactorial nature of this disorder, potentially paving the way for the discovery of new biomarkers and/or disease-modifying treatments.

Two of the most commonly used rodent models in AD research are 3xTg and 5xFAD mice^22,23^. Through the overexpression of familial AD-associated genes, these models collectively recapitulate several key pathological features, including the formation of amyloid-beta (Aβ) plaques, neurofibrillary tangles, neuroinflammation, and cognitive deficits. However, a comprehensive metabolic phenotyping and multi-organ characterization of the 3xTg and 5xFAD mouse models had yet to be conducted. Here, we generate and share a large centralized untargeted metabolomics dataset comprising 2,266 unique biological samples – including brain, serum, liver, cecal content, and feces – from 307 animals (C57BL/6J, 3xTg, and 5xFAD) with or without microbial colonization (specific pathogen free [SPF] or germ-free [GF]) and with additional paired fecal metagenomics data from the SPF cohorts. Via machine learning models^24,25^ and molecular networking^26^, we identify cross-tissue molecular dysregulation of carnitines, bile acids, B vitamins, *N*-acyl lipids, indoles, polyamines, and many other molecules, suggesting a systemic increase of oxidative stress in AD via mitochondrial dysfunction. Further, we find microbially-modulated carnitines as major discriminatory features in the serum of 3xTg animals, including benzoyl-, phenylacetyl- and phenylpropionyl-carnitine. Remarkably, we mapped phenylacetyl-carnitine to multiple human AD cohorts, where this metabolite was positively associated with aging and cognitive impairment. Additionally, 3xTg animals were characterized by depletion of *Akkermansia muciniphila* and an enrichment of *Mucispirillum schaedleri*, *Helicobacter*, *Desulfovibrio*, *Escherichia*, and *Shigella* species, which correlated with serum levels of phenylacetyl-carnitine. Finally, we build and introduce tissueMASST, an open-source mass spectrometry (MS) search tool to investigate known or unknown MS/MS spectra across public metabolomics repositories, facilitating contextualization and interpretation of molecular findings in animal studies to human metabolomics datasets, with associated information on tissue localization and disease status.

This resource represents one of the largest publicly available MS datasets on AD animal models and, together with its associated tools, enables extensive future investigation of uncharacterized molecules and metabolic interactions across multiple tissues in the context of AD.

## RESULTS

### Metabolomics Atlas, Metagenomics, and Tools Development

The metabolomics profiles from SPF and GF cohorts of 3xTg and 5xFAD mice were compared with those of C57BL/6J wild-type (WT) controls. Two separate studies were conducted (**Figure 1A**). In the 3xTg study, animals were euthanized at 7, 12, and 15 months of age and at each time point brain, serum, liver, cecal content, and feces were harvested. In the 5xFAD study, animals were euthanized at 5 and 8 months of age with the same tissues collected. These samples constituted the Sacrifice cohort (n=1,524). Additionally, feces were sampled monthly from the cages of the SPF groups to generate the Longitudinal cohort (n=747). Importantly, both sexes were investigated using a balanced study design (**Figure S1**). Untargeted metabolomics data were acquired using ultra high performance liquid chromatography followed by tandem mass spectrometry (UHPLC-MS/MS) and used to generate a comprehensive and explorable cross-tissue molecular network. Additionally, a complementary MS search tool, tissueMASST, was developed to bridge information obtained from animal studies to human datasets. In total, matching tissue samples were obtained for 307 distinct rodents (**Figure 1B**). Microbial DNA was co-extracted from the fecal samples of SPF animals via the Matrix Method^27^, facilitating the generation of directly matching shallow shotgun metagenomics sequencing data (n=666). Untargeted metabolomics and metagenomics data, along with associated metadata, are publicly available via [GNPS/MassIVE](https://massive.ucsd.edu/ProteoSAFe/static/massive.jsp) and [Qiita](https://qiita-rc.ucsd.edu/). The dataset represents one of the largest publicly available centralized multi-omics resources for AD murine models and greatly expands previous untargeted mouse metabolomics data, with associated metadata, by 25% for serum samples and up to 58% for cecal samples (**Figure 1C**).

**Figure 1.**
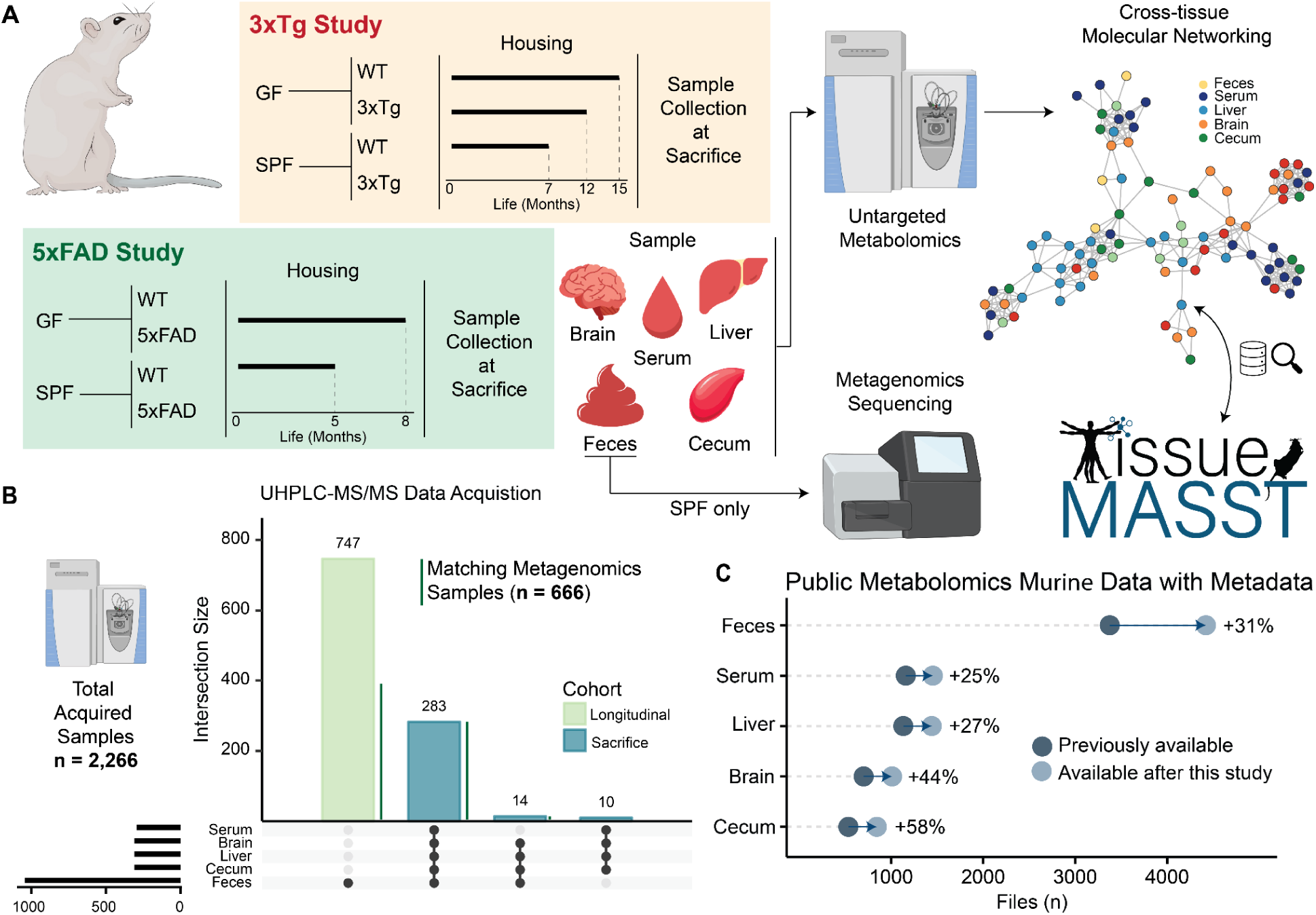
Study Design and Expansion of Public Untargeted Metabolomics Data for Mouse Models. **(A)** Brain, serum, liver, cecal content, and feces were collected at sacrifice from two different studies investigating genetic AD mice (3xTg and 5xFAD) with or without microbial colonization and generated the Sacrifice cohort. Fecal samples were also collected on a monthly basis from SPF mice and constitute the Longitudinal cohort. Samples were analyzed via untargeted UHPLC-MS/MS and shallow shotgun metagenomics sequencing. Metabolomics features were traced across the different tissues to generate a cross-tissue molecular network and tissueMASST, a companion MS/MS search tool described in **Figure 4**, was developed to contextualize animal metabolomics findings in public human data. **(B)** Upset plot of analyzed samples via UHPLC-MS/MS (n=2,266). Matching samples were collected from 307 animals. Paired metagenomics data were acquired from 666 fecal samples. An equal number of female and male animals were investigated in both studies (**Figure S1**). **(C)** Expansion of public murine MS/MS data availability with associated metadata across the 3 major metabolomics repositories ([GNPS/MassIVE](https://massive.ucsd.edu/ProteoSAFe/static/massive.jsp?redirect=auth), [MetaboLights](https://www.ebi.ac.uk/metabolights/), and [Metabolomics Workbench](https://www.metabolomicsworkbench.org/)). Artworks obtained from Canva and BioRender.

### 3xTg and 5xFAD Mice Have Distinct Metabolic Profiles

Clustering of the untargeted metabolomics data was observed across studies (3xTg vs 5xFAD) and cohorts (Sacrifice vs Longitudinal) for each tissue type via principal component analysis (PCA) of the robust center log ratio (RCLR) transformed extracted peak areas (**Figure S2**). Accordingly, downstream data analysis was performed after stratification. Molecular features affected by genotype were extracted via partial least square discriminant analysis (PLS-DA) models from animals on SPF or GF backgrounds (*e.g.*, 3xTg_SPF vs WT_SPF or 3xTg_GF vs WT_GF). Stratifications by genotype and colonization were used to identify molecular features affected by genotype, colonization, or both (**Figure S3**). PLS-DA models displayed superior overall performance, measured as lower classification error rate (CER), in 3xTg animals compared to 5xFAD mice (**Figure 2A**). Interestingly, cecal and brain metabolic profiles returned the best classification performances for genotype in both studies. Lists of extracted metabolic features of interest, with variable importance (VIP) score > 1, differentiating genotypes in mice with a microbiome (*e.g.*, 3xTg_SPF vs WT_SPF) for each tissue type in the two studies are available in **Tables S1-S12**. Additionally, these tables contain information on the influence of colonization or genotype on each individual feature obtained via the stratification workflow described in **Figure S3**. In the 3xTg study, discriminating features were influenced in a similar fashion by genotype (∼25% of total), colonization (∼8%), or both (∼3%) across brain, serum, and liver, while ∼20% were influenced by colonization, ∼10% by genotype, and ∼3% by both for the cecum (**Figure 2B**). Natural log ratios of the features associated with WT (numerator) or AD (denominator) obtained from the PLS-DA models successfully discriminate genotypes across all tissues in both studies (**Figure 2C**). Carnitines, bile acids, B vitamins, N-acyl lipids, indoles, polyamines, and lysophosphatidylcholines (lysoPCs) were among the most discriminating annotated molecules. In the 3xTg study, host age or sex did not alter genotype discriminating power (**Figure S4A**), although male transgenic animals presented exacerbated ratios compared to females in liver, cecal, and fecal profiles (**Figure S4B**). In the 5xFAD study, sex stratification highlighted that ratios could still separate AD animals from controls in females, but separation was lost for brain and liver profiles in male mice (**Figure S5**). For the rest of this study, we focus on in-depth profiling of 3xTg mice to identify dysregulated metabolic pathways of interest and uncharacterized molecules modulated by the gut microbiome, though all data from 5xFAD mice are publicly available as described.

**Figure 2.**
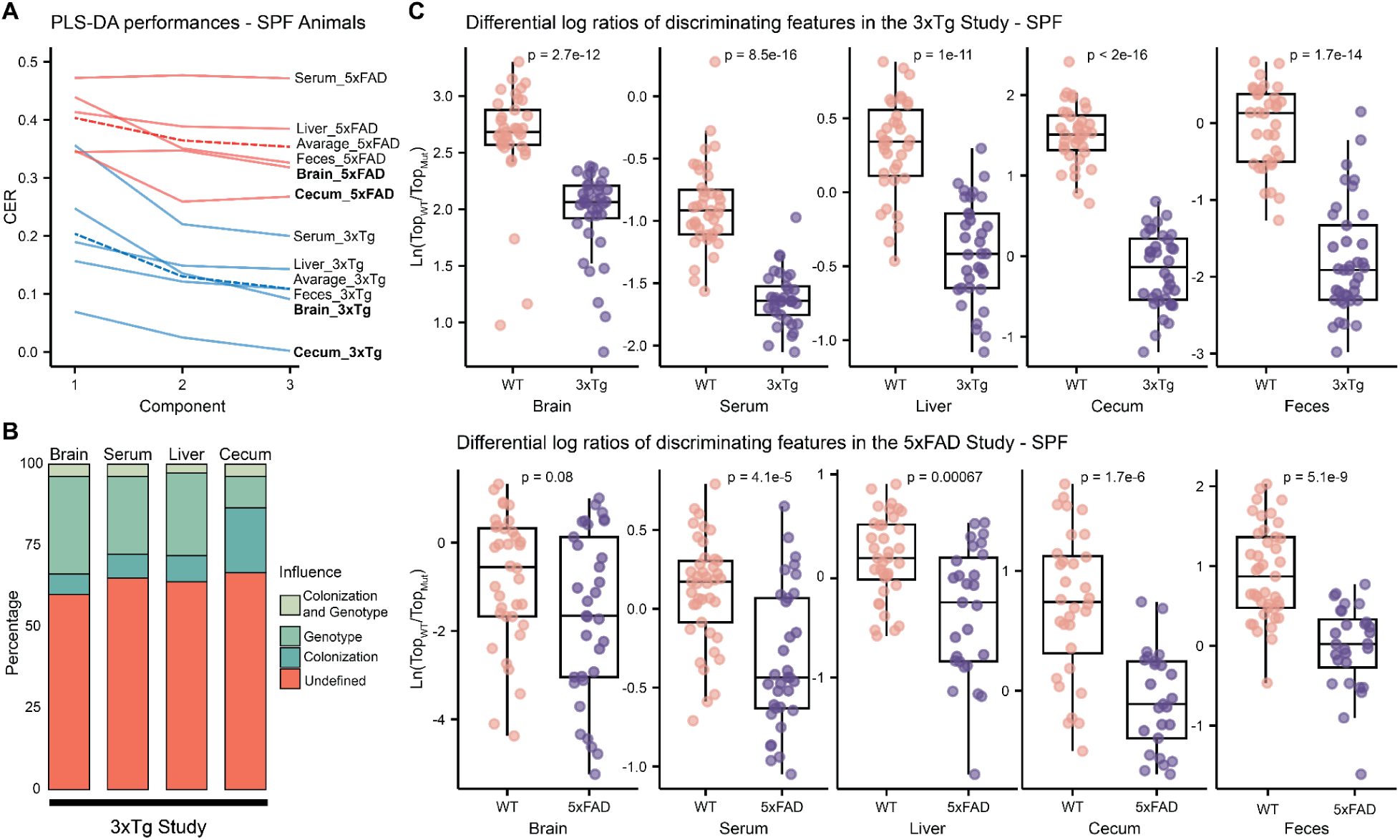
Genotype and Microbiome Status Influence Tissue Metabolic Profiles. **(A)** PLS-DA model performances, per tissue, for genotype classification in SPF animals from the 3xTg and 5xFAD studies. Dashed lines represent average CER across tissue per study. **(B)** Influence of colonization or genotype on discriminating features of interest extracted from PLS-DA models obtained via the nested workflow described in **Figure S3**. A stronger colonization effect was observed in cecum. **(C)** Natural log ratios of significant features decreased (numerator) or increased (denominator) in AD animals, compared to WT, extracted from the respective PLS-DA models constructed on SPF rodents only. Significance tested via Wilcoxon test. Boxplots show first (lower) quartile, median, and third (upper) quartile.

### Genotype-driven Metabolic Changes in 3xTg Mice Indicate Increased Systemic Oxidative Stress

Thousands of metabolic features differentiate molecular profiles across tissues between WT and AD models (**Tables S1-S12**) suggesting systemic dysregulation across several classes of molecules (**Figure 3**), including bile acids, B vitamins, N-acyl lipids, neurotransmitters, carnitines (**Figure 4**), and many others.

**Figure 3.**
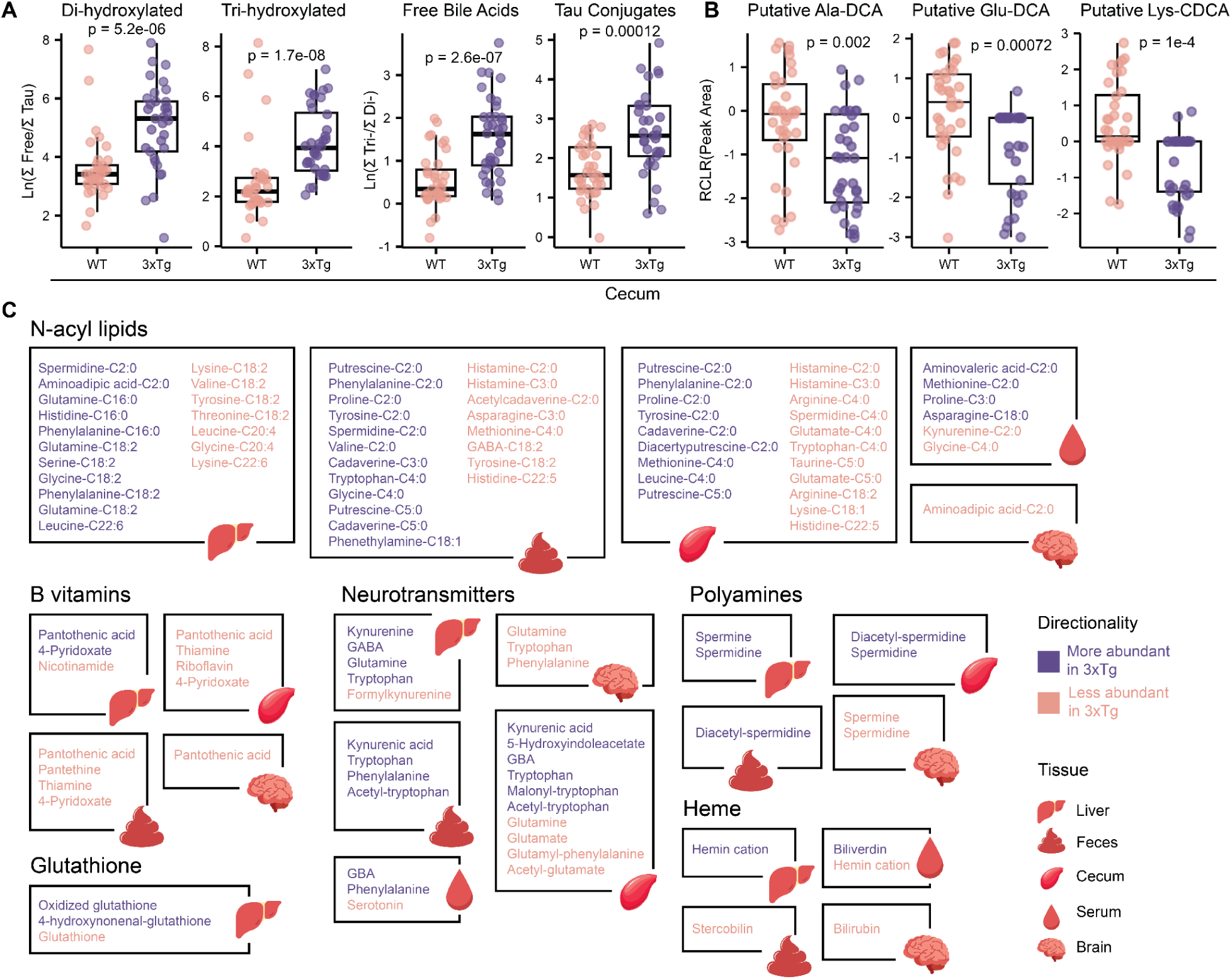
Annotated Molecular Features Dysregulated in 3xTg Mice with an SPF Microbiome. **(A)** Natural log ratios of free over taurine conjugated di- and tri-hydroxylated bile acids and of tri- over di- free or taurine conjugated bile acids in the cecum. Significance tested via Wilcoxon test. **(B)** Univariate analysis of putative microbial di-hydroxylated bile acids conjugated to amino acids (Ala, alanine; Glu, glutamate; Lys, lysine) in the cecum. Significance tested via Wilcoxon test. **(C)** Schematic representation of molecules altered in 3xTg mice on a SPF background found via PLS-DA models. Affected classes included N-acyl lipids, B vitamins, neurotransmitters, ethanolamides, and polyamines and lysoPCs among the others. Boxplots show first (lower) quartile, median, and third (upper) quartile. Annotations are based on MS/MS spectral matching, which correspond to a level 2 or 3 annotation according to the Metabolomics Standard Initiative (MSI)^52^.

**Figure 4.**
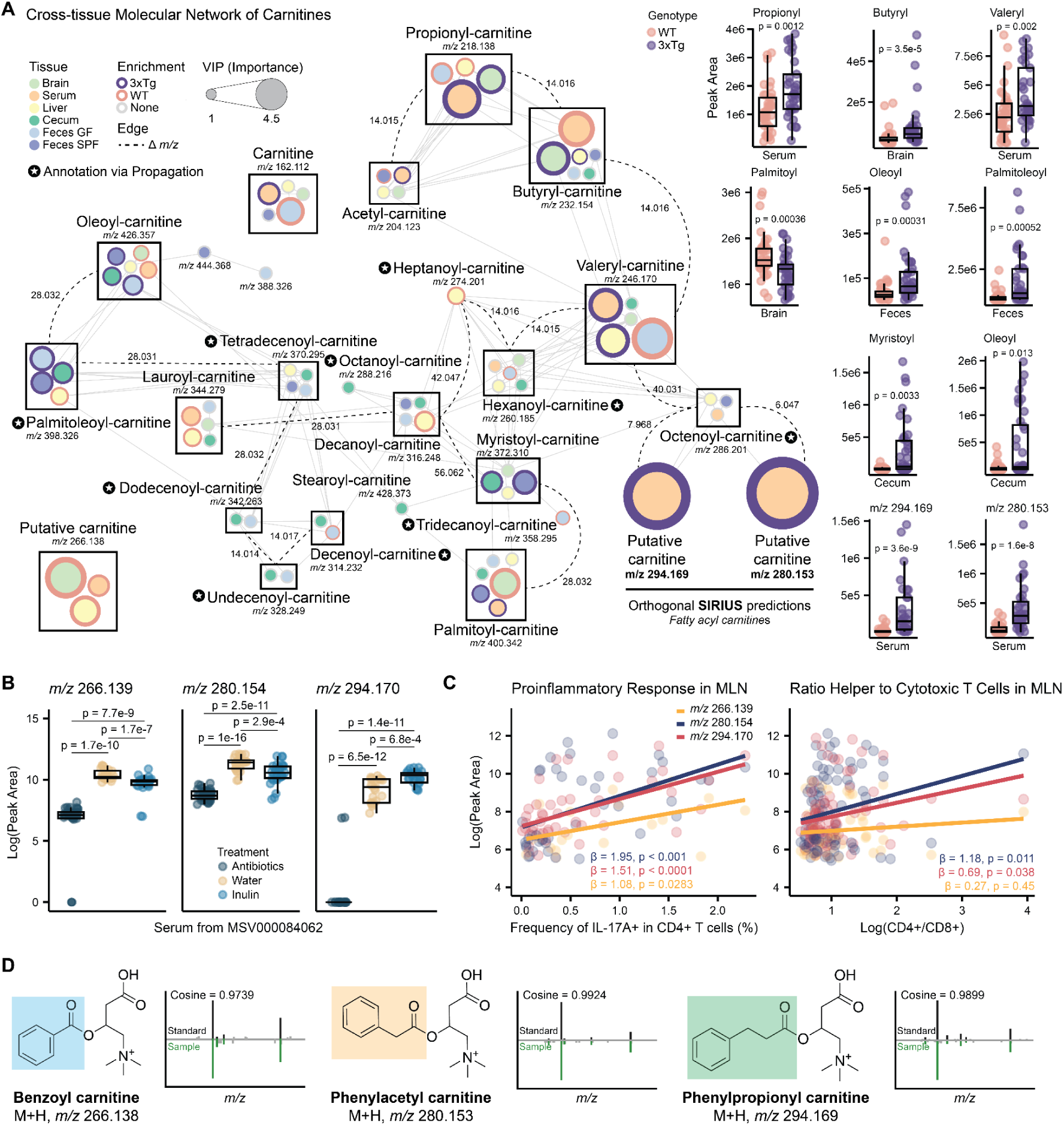
Carnitine Dysregulation Across Multiple Tissues in 3xTg Animals. **(A)** Cross-tissue molecular network of carnitines in the 3xTg study. Each node represents a molecule and is color-coded based on tissue type. Same molecules from different tissues are boxed together and edges, dashed lines, report the delta *m/z* between molecules. Molecules enriched in either WT or 3xTg animals are reported via colored labels and nodes are scaled based on the VIP scores extracted from the per tissue PLS-DA models. Annotations were propagated through the network. Boxplots of selected carnitines of interest showcase genotype differences via univariate analyses. **(B)** Extracted carnitines’ peak areas detected in serum from an external murine dataset ([MSV000084062](https://massive.ucsd.edu/ProteoSAFe/dataset.jsp?task=abbb4996c26843f9a8c73eba0d743794)). Antibiotics treatment significantly decreased the relative abundance of the putative carnitines. Pairwise Wilcoxon test followed the Benjamini-Hochberg (BH) correction. Boxplots show first (lower) quartile, median, and third (upper) quartile. **(C)** Linear correlations between carnitines’ peak areas and IL-17A+ CD4+ T cells and CD4+/CD8+ ratio in the mesenteric lymph nodes (MLN). Linear models accounted for sex as a covariate. The molecules *m/z* 280.153 displayed the strongest correlation to the inflammatory markers. **(D)** MS/MS spectral matching of synthesized standard for benzoyl-, phenylacetyl-, and phenylpropionyl-carnitine.

3xTg animals presented lower levels of taurine conjugated di- and tri-hydroxylated bile acids, including putative taurodeoxycholic (TDCA) and taurocholic acid (TCA), in the serum, liver, and cecum; less di-hydroxylated bile acids, such as putative deoxycholic (DCA), chenodeoxycholic (CDCA), and ketodeoxycholic acid (ketoDCA), in the cecum and feces; and lower levels of putative cholic acid (CA) in serum, liver, and cecum, together with less putative mono-hydroxylated lithocholic acid (LCA) in the cecum. Importantly, bile acids act on both the farnesoid X receptors (FXR) and G protein-coupled receptors TGR5^28^, which regulate oxidative stress^29,30^. Ratios in the cecum of free tri- to di-hydroxylated bile acids and their taurine conjugates also showed significant separation based on genotype (**Figure 3A**). Additionally, microbial bile acids, such as putative Ala-DCA, Glu-DCA, and Lys-CDCA were observed in higher abundances in the cecum of WT animals (**Figure 3B**), possibly due to reduced microbial activity or substrate (TCA) availability.

The recently described N-acyl lipids^31^, including many derived from short chain fatty acids (C2 to C5), were also dysregulated in AD mice (**Figure 3C**). The cecum and feces of 3xTg animals were depleted of histamine-C2:0 and -C3:0 while enriched in polyamines and aromatic amino acids conjugates of C2 to C5, including putrescine, cadaverine, spermidine, phenylalanine, tyrosine, tryptophan, and proline. Interestingly, while feces and cecum showed differences based on short chain (C2-C5) conjugates, differential abundances in the liver were driven by long chain (C16-C22) conjugates. Notably, none of these molecules have been previously associated with aging or AD in humans or their associated mouse models and their biological roles are currently undefined.

B vitamins, which are involved in fatty acid metabolism via β-oxidation^32–34^, were generally depleted in 3xTg animals across multiple tissues (**Figure 3C**). Pantothenic acid (vitamin B5), rate-limiting precursor of Coenzyme A (CoA), which is essential in the β-oxidation process^35^, was decreased in the brain, cecum, and feces of 3xTg mice together with thiamine (vitamin B1) in the cecum and feces, riboflavin (vitamin B2) in the cecum and 4-pyridoxate, product of vitamin B6 catabolism, in the cecum and feces. Interestingly, increased vitamin B5 and 4-pyridoxate and decreased nicotinamide, water soluble form of vitamin B3, were detected in the liver of AD animals compared to WT mice. Importantly, the liver of 3xTg animals also presented higher levels of oxidized glutathione (GSSG) and 4-hydroxynonenal-glutathione (HNE-GSH) and lower levels of glutathione (GSH). GSH has a crucial role in β-oxidation protecting from reactive oxygen species (ROS)-mediated damage generated during fatty acid breakdown^36^ and higher abundances of GSSG and HNE-GSH are reflective of increased oxidative stress due to mitochondrial dysfunction^37^. Notably, GSSG can be reduced back to GSH at the expense of NADPH, whose precursor vitamin B3^38^, was depleted in the liver of 3xTg mice. Hemin, known to be involved in mitochondrial dysfunction^39^, was more abundant in the liver of 3xTg mice, but lower in the serum. Biliverdin was enriched in the serum of AD animals, while bilirubin and stercobilin were depleted in the brain and feces respectively. Bilirubin is known to protect from neuronal damage in the brain by scavenging for ROS^40^ and higher levels of biliverdin in serum can be indicative of increased presence of ROS^41^.

Neurotransmitters were also dysregulated in 3xTg mice (**Figure 3C**). Tryptophan metabolism appeared to be channeled into the kynurenine pathway, known to promote oxidative stress^42^, as kynurenine and kynurenic acid were higher in the liver, cecum, and feces of AD mice, together with tryptophan itself. Additionally, 3xTg animals displayed lower levels of serotonin in the serum and higher levels of 5-hydroxyindoleacetate (5-HIAA), a primary metabolite of serotonin, in the cecum. The γ-aminobutyric acid (GABA) shunt pathway was also dysregulated, with 3xTg mice showing increased GABA in the liver and more 4-guanidinobutanoate (GBA), a metabolite of GABA, in the serum and cecum. Serotonin and GABA have been reported to mitigate oxidative stress^43,44^. Glutamate and glutamine were also depleted in the cecum of 3xTg mice while glutamine was higher in the liver but lower in the brain. The brains of 3xTg animals also presented lower levels of tryptophan and phenylalanine, which was also enriched in the serum and feces of AD animals. Importantly, the polyamines spermine and spermidine were more abundant in the liver of 3xTg mice, but depleted in the brain. These molecules are known to prevent neurodegeneration in the brain by promoting mitophagy^45,46^, a form of autophagy that clears damaged mitochondria, while in the liver they are considered biomarkers for hepatocytes regeneration^47^.

Several other annotated molecules were found to be dysregulated (**Figure S6**). These included 1) ethanolamides, which are produced in response to cellular injury^48,49^; 2) ornithine, which can stimulate ROS and GABA production by the astrocytes^50^; 3) hippurate, which induces mitochondrial ROS production^51^, 4) purine and pyrimidine derivatives, 5) lysoPCs, 6) linoleic acid metabolites, and hundreds of additional related uncharacterized molecules.

### Cross-Tissue Molecular Network Reveals Dysregulation of Carnitines in 3xTg Mice

To track metabolic features across multiple tissues, we created an interactive cross-tissue molecular network - downloadable via GitHub - by networking tissue-specific consensus MS/MS spectra generated in this study. The cross-tissue molecular network generated from the 3xTg study revealed that carnitine metabolism was broadly disrupted across multiple tissues (**Figure 4A**). Compared to WT, 3xTg mice presented higher levels of acetyl-carnitine in serum, propionyl-carnitine in the brain and serum, butyryl-carnitine in the brain and liver, valeryl-carnitine in serum and liver, myristoyl-carnitine and oleoyl-carnitine in the cecum and feces, stearoyl-carnitine and arachidonoyl-carnitine in the liver, and palmitoyl-carnitine in serum and feces. Conversely, 3xTg mice had lower levels of acetyl-carnitine in feces, propionyl-carnitine in the liver, butyryl-carnitine in serum, oleoyl-carnitine in the brain and serum, palmitoyl-carnitine and arachidonoyl-carnitine in the brain, lauroyl-carnitine in serum and liver, and decanoyl-carnitine in the liver. Additionally, propagating information through the cross-tissue molecular network allowed us to annotate palmitoleoyl-carnitine, which was enriched in the cecum and feces but depleted in the liver of 3xTg mice. Interestingly, three unannotated spectra (*m/z* 266.138, 280.153, and 294.169), two of which were more abundant in the serum of 3xTg animals and one in serum, brain, and liver of WT, were identified as possible carnitines, as they were in the carnitine subnetwork and putatively classified as fatty-acyl carnitines via SIRIUS^53,54^ (**Figure 4A**). Investigating the colonization effect on these molecules via GF animals and the stratification workflow suggested that they are microbially-modulated, either produced by microbes or co-metabolized by pathways in microbes and mice (**Table S2**). To confirm this hypothesis, an external untargeted metabolomics dataset acquired from serum of C57BL/6J mice exposed either to regular drinking water, a cocktail of antibiotics, or inulin was reanalyzed. Interestingly, the abundance of these molecules was reduced in response to antibiotic exposure, while opposing trends were observed for inulin, a prebiotic that fosters the growth of many gut bacteria (**Figure 4B**). Furthermore, immunological data co-acquired from the 3xTg animals at 12 months of age, presented in the companion manuscript by Bostick et al.^55^ profiling the immune response, highlighted a positive correlation between two molecules (*m/z* 280.153 and 294.169) and the IL-17A+ CD4+ T cell population (T helper 17 cells; Th17) and the CD4+/CD8+ ratio in the mesenteric lymph nodes, irrespective of sex (**Figure 4C**).

Given the likely microbial nature of these molecules and their association to inflammation in an AD mouse model, we proceeded with structural elucidation after hypothesizing the unannotated metabolites to be carnitine conjugates of the microbially-derived benzoate, phenylacetic acid, and phenylpropionic acid. We subsequently synthesized molecular standards and proceeded to Level 1 annotation via MS/MS spectra and retention time matching (**Figure 4D**). The spectra of benzoyl-carnitine (M+H, *m/z* 266.138), phenylacetyl-carnitine (M+H, *m/z* 280.153), and phenylpropionyl-carnitine (M+H, *m/z* 294.169) are now publicly available via the GNPS library to facilitate annotation in other untargeted metabolomics datasets.

### Fecal Microbial Profiles are Associated with Serum Phenylacetyl-carnitine in 3xTg Mice

Fecal samples collected from SPF (colonized) animals in the Sacrifice and Longitudinal cohorts were analyzed via shotgun metagenomics. Beta diversity analysis via phylogenetic Robust Aitchison PCA (phylo-RPCA) was used to explore community differences based on metadata information. Similar to the untargeted metabolomics analyses, a cohort effect was observed in both the 3xTg and 5xFAD studies (**Figure S7A**). Focusing on the Sacrifice cohorts, a significant genotype effect (PERMANOVA, pseudo-F = 28, p < 0.001) was observed in the 3xTg study (**Figure 5A**) but not in the 5xFAD study (**Figure S7B**; PERMANOVA, pseudo-F = 0.14, p = 0.885). Conversely, a small sex effect (PERMANOVA, pseudo-F = 3.9, p = 0.026) was observed in the 5xFAD study but not in the 3xTg one (**Figure S7C**). No significant alpha diversity differences were detected based on genotype (**Figure S7D**). Log ratios using the top and bottom features associated with the relevant principal components from the phylo-RPCAs were generated. In the 3xTg study, the second component of the phylo-RPCA was used, as it separated samples based on genotype (**Figure 5A**). Selected log ratios, including *Helicobacter bilis*, *Mucispirillum schaedleri,* and *Akkermansia muciniphila* displayed significant separation between WT and 3xTg mice (**Figure 5B**). The selected log ratios from the Sacrifice cohort were then applied to the Longitudinal cohort (**Figure 5C**), which displayed a significant separation based on genotype during early-life that converged later in life (linear mixed effect model, β = -8.46, p = 3.37e-10). A similar strategy was used to create log ratios in the 5xFAD study, but no significant differences were observed (**Figure S7E**).

**Figure 5.**
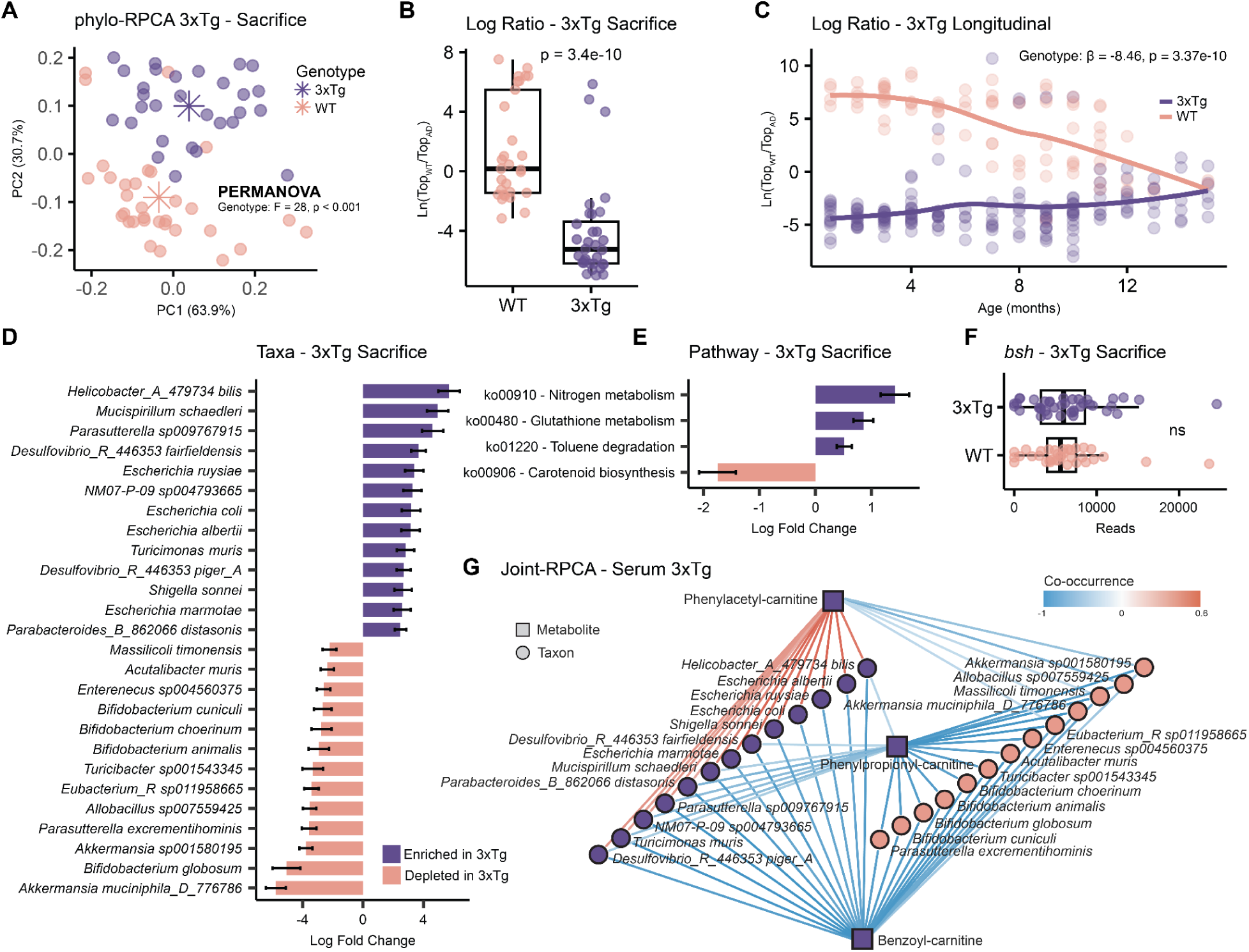
3xTg Mice Display Distinct Fecal Microbial Profiles that Correlate with Serum Carnitines. (**A**) phylo-RPCA of fecal samples collected in the 3xTg study from the Sacrifice cohort showed a significant separation based on genotype (PERMANOVA, pseudo-F = 28, p < 0.001). (**B**) Natural log ratios of the top/bottom ranked OGUs from Axis 2 of phylo-RPCA of 3xTg Sacrifice cohort. Significance determined by two-sided Mann-Whitney U test. (**C**) Log ratio defined via Sacrifice cohort was then applied to the Longitudinal cohort samples. A strong separation was observed during the first months of life, which then converged as mice aged. Linear mixed effect model used to account for repeated measures of fecal samples collected from the same cage. (**D**) Differential abundance analysis via ANCOM-BC2 of OGUs collapsed to species level of 3xTg Sacrifice cohort (ANCOM-BC2, LFC > 2, p adjusted < 0.05). AD animals were enriched in *M. schaedleri* and *Helicobacter*, *Desulfovibrio*, *Escherichia*, and *Shigella* related species and depleted in *A. muciniphila*. (E) Differential abundance analysis via ANCOM-BC2 of KEGG pathways. 3xTg animals showed enrichment in pathways related to nitrogen, glutathione, and toluene metabolism. (**F**) No differences in encoded *bsh* gene levels between WT and 3xTg animals were observed in the 3xTg study. Significance tested via Wilcoxon test. (**G**) Network of bacteria-metabolites co-occurrences in serum obtained via joint-RPCA. Phenylacetyl-carnitine, which was enriched in the serum of 3xTg animals, showed positive co-occurrence with fecal bacteria enriched in 3xTg mice. On the contrary, phenylpropionyl-carnitine, also enriched in 3xTg serum, displayed negative co-occurrence with all taxa of interest. Only edges with absolute co-occurrence > 0.3 are reported.

Differential abundance analysis was also conducted using ANCOM-BC2 at species level. 3xTg mice harbored increased abundance of *M. schaedleri* and various species belonging to the *Helicobacter*, *Desulfovibrio*, *Escherichia*, and *Shigella* genera and decreased abundance of *A. muciniphila* strains and *Bifidobacterium* species (**Figure 5D**). In the 5xFAD study, only one species belonging to the *CAG-286* genus appeared to be enriched in this AD mouse model compared to WT (**Figure S7F**). KEGG pathways analysis of the 3xTg Sacrifice cohort samples identified an enrichment in nitrogen metabolism, glutathione metabolism, and toluene degradation and a depletion of carotenoid biosynthesis in 3xTg animals (**Figure 5E**), pathways also partially affected in the 5xFAD mice (**Figure S7G**). Given the observed differences in the microbial carnitines related to phenylalanine metabolism^56^ and in bile acids conjugation, selective gene searches for *fldC* (phenyllactyl-CoA dehydratase subunit) and *bsh* (bile salt hydrolase/transferase) were conducted using DIAMOND^57^. The *fldC* gene was largely absent in the analyzed samples (**Figure S7H**). No difference in encoded *bsh* gene levels were observed between WT and 3xTg animals (**Figure 5F**), suggesting that bile acid differences might be attributed to a reduction in taurine conjugates production in the liver rather than microbial activity.

Finally, multi-omics data integration was performed via joint-RPCA^25,58^. While the co-occurrence of all microbes and metabolites that passed thresholds were calculated for each tissue (**Table S13-S17**), a network focused on differential fecal taxa and serum carnitines of interest was generated (**Figure 5G**). Interestingly, phenylacetyl-carnitine positively and negatively co-occurred with microbial species that were observed to be enriched or depleted in the 3xTg animals, respectively, compared to WT mice. Conversely, benzoyl- and phenylpropionyl-carnitine negatively co-occurred with bacterial species altered in this mouse model. These findings suggest that microbiome dysbiosis in 3xTg mice may contribute to alterations in serum levels of carnitines that were also dysregulated in this model.

### Phenylacetyl-carnitine is Associated with Aging and Cognitive Impairment in Humans

To leverage the data generated in this preclinical study and contextualize it across human metabolomics datasets, we developed tissueMASST, a search tool allowing querying for known and unknown MS/MS spectra against public tandem mass spectrometry data acquired from human and rodent samples (**Figure 6A**). tissueMASST improves on previous MS/MS search tools^59–61^ by incorporating MS/MS data from all three major public metabolomics repositories, [GNPS/MassIVE](https://massive.ucsd.edu/ProteoSAFe/static/massive.jsp), [MetaboLights](https://www.ebi.ac.uk/metabolights/)^62^ and [Metabolomics Workbench](https://www.metabolomicsworkbench.org/)^63^. Additionally, this tool unlocks access to tissue-associated metadata, when available, such as disease status, sex, and age. The reference database of tissueMASST comprises 44,992 public MS/MS files as of April 2025. Of these, 27,954 were acquired from human tissues, 11,422 from mice, and 1,328 from rats. tissueMASST encompasses data from 58 different tissues, including the brain, cerebrospinal fluid, feces, heart, liver, lung, milk, plasma, serum, and several parts of the gastrointestinal tract, and more than 20 different diseases, such as AD, type 2 diabetes (T2D), inflammatory bowel disease (IBD), acquired immunodeficiency syndrome (AIDS), and others (**Figure S8**). tissueMASST is available as a standalone web application (https://masst.gnps2.org/tissuemasst/), where users can search single MS/MS spectra via an intuitive interface, or as Python script, which allows for batch searches of multiple spectra and the co-search against other domain MASSTs, such as microbeMASST^60^, plantMASST^61^, and foodMASST^64^; enabling the investigation of possible bio-localization and origin of molecules of interest.

**Figure 6.**
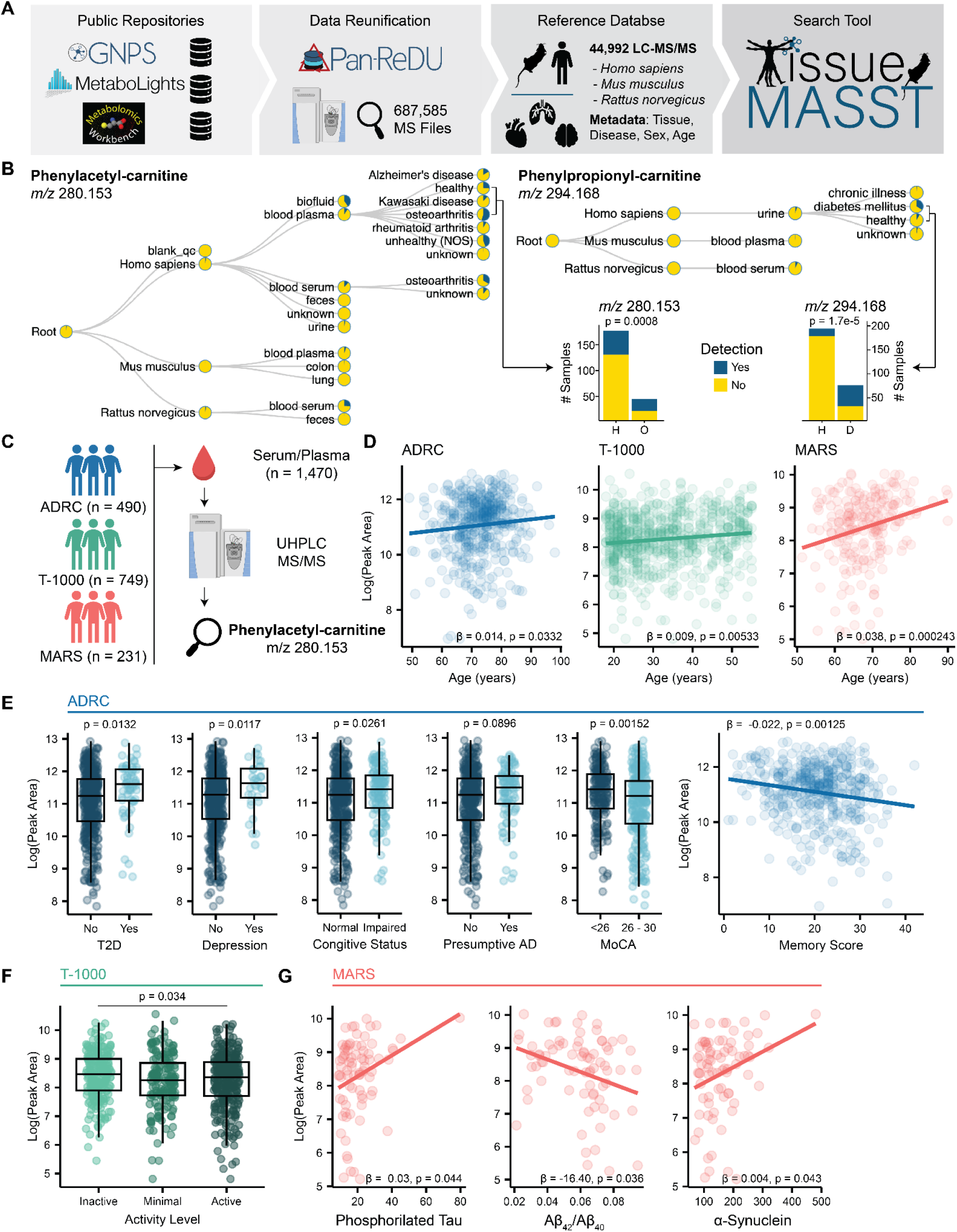
Serum Phenylacetyl-carnitine Levels Correlates with Aging and Cognitive Impairment. **(A)** tissueMASST leverages the 3 major public metabolomics data repositories ([GNPS/MassIVE](https://massive.ucsd.edu/ProteoSAFe/static/massive.jsp), [MetaboLights](https://www.ebi.ac.uk/metabolights/), and [Metabolomics Workbench](https://www.metabolomicsworkbench.org/)). Metadata reunification via [PanReDU](https://redu.gnps2.org/selection/)^69^ across repositories listed 687,585 files. Of these, 44,992 MS/MS files were from *Homo sapiens*, *Mus musculus*, and *Rattus norvegicus*. Metadata associated with these files, such as biolocalization (tissue), disease status, sex, and age, was extracted and organized. **(B)** tissueMASST search outputs. Phenylacetyl-carnitine (PAC, *m/z* 280.153) was mainly detected in human serum and blood samples and appeared to be more prevalent in patients with osteoarthritis compared to healthy subjects. Phenylpropionyl-carnitine (*m/z* 294.169) was mainly found in human urine and is enriched in subjects with type II diabetes (T2D) compared to healthy individuals. Statistical difference was calculated via Chi-squared test. **(C)** PAC was detected in plasma and serum datasets, including the ADRC, T-1000, and MARS cohorts (combined n=1,470). Datasets were reprocessed with MZmine to extract PAC peak areas. **(D)** Concentration of PAC positively correlated with age across datasets. Linear model adjusted for sex and BMI. **(E)** PAC was observed in higher abundance in subjects with T2D, depression, and subjects with putative cognitive impairment. MoCA scores were adjusted for education. When looking at medical diagnosis, subjects with MCI or dementia presented higher serum PAC, while a tendency was observed in subjects with MCI or dementia and a presumptive AD diagnosis. Finally, PAC negatively correlated with Craft Story- Delayed Recall Verbatim scores. P values were corrected for age, sex and BMI. The T2D model was also corrected for hypercholesterolemia. **(F)** Active individuals in the T-1000 cohort presented lower levels of PAC. **(G)** PAC serum levels in the MARS cohort correlated with CSF levels of phosphorylated tau protein, α-synuclein, and Aβ_42_/Aβ_40_ ratio.

Searching phenylacetyl-carnitine (PAC) and phenylpropionyl-carnitine (PPC) via tissueMASST returned results mapping to human blood and urine (**Figure 6B**). PAC appeared to be enriched in inflammatory conditions and, importantly, tissueMASST was able to match PAC to several human AD datasets where metadata is available through the [AD Knowledge Portal](https://adknowledgeportal.synapse.org/), encompassing hundreds of serum and plasma samples generated by the [Alzheimer Gut Microbiome Project](https://alzheimergut.org/) (AGMP) Consortium and [Accelerating Medicines Partnership for Alzheimer Disease](https://fnih.org/our-programs/accelerating-medicines-partnership-amp/amp-alzheimers-disease-2-0/) (AMP-AD). The datasets of interest were the Alzheimer’s Disease Research Center (ADRC), the Tulsa-1000^65,66^ (T-100), and the Microbiome in Alzheimer’s Risk Study^67^ (MARS). Study population descriptions are provided in **Methods**. We reprocessed untargeted metabolomics data acquired from the ADRC (n=490), T-1000 (n=749), and MARS (n=231) cohorts to extract the peak areas of PAC (**Figure 6C**) and confirmed a Level 1 annotation (**Figure S9**). Remarkably, PAC was positively correlated with aging, after adjusting for sex and BMI, in all three datasets (**Figure 6D**). In the ADRC cohort, PAC was enriched in subjects with T2D, even after adjusting for hypercholesterolemia, or depression (**Figure 6E**). Interestingly, subjects with any form of cognitive impairment (e.g., clinical diagnosis of either mild cognitive impairment [MCI] or dementia) had higher levels of PAC in their blood than unimpaired individuals, again after adjusting for sex, age, and BMI (**Figure 6E**). A similar trend was observed for subjects with AD-specific cognitive impairment, although significance was lost after adjusting for age, sex, and BMI (p = 0.0896). In terms of cognitive abilities, participants who screened as impaired on the Montreal Cognitive Assessment^68^ (i.e., score <26) had higher PAC levels. Finally, PAC was associated with poorer performance on a test of memory, which is the cognitive domain most impacted by AD (**Figure 6E**). In the T-1000 cohort, which focuses on major depressive disorders in a younger population, no diagnostic information was observed. However, serum PAC was more abundant in sedentary subjects (**Figure 6F**). Finally, in the MARS cohort, cerebral spinal fluid biomarkers were obtained from a subset of 77 individuals and PAC was positively correlated to phosphorylated tau protein and α-synuclein levels and negatively correlated to Aβ_42_/Aβ_40_ ratio after adjusting for age, sex, and BMI (**Figure 6G**). Taken together, these findings suggest a relevant role for phenylacetyl-carnitine in aging and cognitive impairment, expanding knowledge for potential contributing interactions between the gut microbiome and AD.

## DISCUSSION

We present a centralized untargeted metabolomics resource of multiple tissues from the 3xTg and 5xFAD animal models of AD, with associated metagenomics data. With more than 2,000 unique biological LC-MS/MS samples acquired from more than 300 animals, with different colonization conditions, genotypes, sex, and age, this resource will enable expansion and acceleration of AD research. Additionally, we release a companion interactive cross-tissue molecular network that can be used to explore unknown molecules in the context of AD murine models. To further facilitate feature tracking across metabolomics datasets, we introduce tissueMASST. This MS/MS search tool allows easy access to public mass spectrometry data acquired from animal models and humans with associated metadata. When used in conjunction with the other domain MASSTs^60,61,64^, researchers can rapidly identify possible bio-localization, disease association, and origin of known and unknown molecules of interest across studies in AD and any other area of biomedical research.

Herein, we deeply investigate previously uncharacterized MS/MS spectra enriched in the serum of 3xTg animals, which we elucidate as benzoyl-carnitine, phenylacetyl-carnitine, and phenylpropionyl-carnitine. Little is known about these molecules, which have not been previously implicated in AD^70–75^. Their precursors, benzoate, phenylacetate, and phenylpropionate likely derive from the microbial metabolism of phenylalanine^56^, but it remains unclear if the conjugation is mediated by microbes or host enzymes. Interestingly, we traced phenylacetyl-carnitine to thousands of human blood samples, including from large clinical AD studies. In these, phenylacetyl-carnitine showed positive correlations with aging and cognitive impairment. Thus, this molecule represents another addition to the list of biologically relevant host-microbial metabolites linked to human conditions. Notably, phenylacetyl-carnitine correlated with the fecal abundance of several microbial taxa. *A. muciniphila*, usually depleted in AD^76^, is known to potentially enhance cognitive function^77–79^ and to reduce inflammation by modulating intestinal macrophages^80^. Interestingly, the dysregulated metabolic state of 3xTg animals suggested the presence of a pro-inflammatory status. Indeed, we observed microbiome-directed inflammation in this model in the companion manuscript^55^. Notably, inflammation results in the activation of macrophages and production of ROS and nitrate^81^, which promotes the growth of *M. schaedleri*^82^. This bacterium is enriched in inflammatory diseases^83^ and resides in the intestinal mucus layer, possibly competing with *A. muciniphila* when the mucus layer thickness is reduced by inflammation^84^. Phenylacetyl-carnitine was also found enriched in subjects with T2D, a condition known to increase AD risk by up to 45–90%^85^ and is usually associated with reduced *A. muciniphila* levels^86^. As T2D and AD are linked to cardiovascular disease^87^, it is interesting to note that phenylacetyl-carnitine has been recently reported as a serum biomarker for ischemic heart disease^88^. Finally, serum phenylacetyl-carnitine was associated with several diagnostic cerebral spinal fluid biomarkers in AD, such as phosphorylated tau protein^89^, α-synuclein^90^, and Aβ_42_/Aβ_40_ ratio^91^. For these reasons, phenylacetyl-carnitine represents a molecule of interest in the context of aging, inflammation, and cognitive impairment and its role should be further explored in relation to metabolic disorders in humans.

Systemic analysis of the untargeted metabolomics data generated across the different tissues also recapitulated key molecular dysregulations previously associated with AD and related to mitochondrial dysfunction and oxidative stress^92–95^. Although carnitines represent a primary indicator of this disruption, several other molecules highlighted by the current study reinforce links to mitochondrial alterations. Dysregulation of B vitamins, essential for β-oxidation^35^, were observed in 3xTg mice. Supplementation of these molecules has been linked to improve cognition in both humans and animal models^96–100^ and vitamin B5 deficiency has been correlated with neuronal damage in AD-associated brain regions^101^. B vitamins are also key regulators of neurotransmitters, such as serotonin, dopamine, and GABA^102^. As reduced levels of serotonin have been observed in AD patients^103^, GABA accumulation in the liver has been mechanistically associated with metabolic dysfunction^104^. The role of the liver in AD pathogenesis, in particular in relation to Aβ clearance and bile acid metabolism, is under active investigation^105,106^. Bile acids, molecules trafficking between and being transformed by the liver and the gut microbiome, have been recently linked to AD with numerous reported associations^11–14,107–111^, with ongoing investigations suggesting the potential use of tauroursodeoxycholic acid in treating AD^112^.

Herein, we highlighted interactions between multiple annotated metabolites and oxidative stress, since this connection is robust and appears relevant to human AD. However, we recognize that hundreds of non-annotated molecules are present and differentially abundant between WT and AD mouse models. The scientific community will be able to investigate pathways of interest via this centralized and publicly-available resource of LC-MS/MS data to discover uncharacterized metabolites, as we showcased for the microbial carnitines in this study. These findings and approaches will further inform studies for mechanistic roles of microbially-modulated metabolites in the context of AD pathogenesis. Finally, the companion tool we developed, tissueMASST, will enable the research community to translate animal model findings to human data, with the goal of accelerating discovery of biomarkers and potential therapeutics.

### Limitations of the study

MS/MS spectra annotation via spectral matching against the public GNPS reference libraries represent a Level 2 or 3 annotation according to the Metabolomics Standards Initiative^52^. Cross-tissue molecular networking is based on MS/MS spectral matching and it is possible that some spectral pairs are missing due to the poor fragmentation or missing fragments during the acquisition of the MS/MS spectra. Germ-free animals have received a cocktail of antibiotics during behavioral testing to maintain their microbiological sterility and this could have potentially altered organ metabolic profiles. Fecal samples in the Longitudinal study were collected randomly from animals in the same cage and they did not have an associated animal ID so investigation of single animal development was not possible. Feature detection across reference datasets present in tissueMASST can be influenced by study-specific extraction methods, workflows, and instrumentation. Thus, lack of detection does not necessarily translate to the molecule being absent in tissues. Differences between experimental cohorts (Longitudinal and Sacrifice) within each study were observed for both microbiome and metabolomics data and can be attributed to differences in sample collection. Longitudinal samples were obtained by collecting freshly produced fecal samples, whereas sacrifice samples were collected from the colon after sacrifice. Finally, 3xTg and WT mice were bred and housed separately before the start of the experiment.

## RESOURCE AVAILABILITY

### Lead contact

Further information and requests for resources should be directed to and will be fulfilled by the lead contacts Pieter C. Dorrestein.

### Materials availability

This study did not generate new unique reagents.

### Data and code availability

- Untargeted metabolomics data and metadata generated for this manuscript are available on [GNPS/MassIVE](https://massive.ucsd.edu/ProteoSAFe/static/massive.jsp) under the following accession codes: [MSV000096035](https://massive.ucsd.edu/ProteoSAFe/dataset.jsp?task=b467af929b9347b8b0aa13f41957fa4d), [MSV000096036](https://massive.ucsd.edu/ProteoSAFe/dataset.jsp?task=8249c40f7e3d4ea8a4e20871725a7ca3), [MSV000096037](https://massive.ucsd.edu/ProteoSAFe/dataset.jsp?task=60905386a3f944e1b859b8d96592980b), [MSV000096038](https://massive.ucsd.edu/ProteoSAFe/dataset.jsp?task=7d99b115a6d346d5b787c7ce10030a79), [MSV000096039](https://massive.ucsd.edu/ProteoSAFe/dataset.jsp?task=55ec7f9e84124ca0827f0d7958192c34), [MSV000093230](https://massive.ucsd.edu/ProteoSAFe/dataset.jsp?task=fbf55f75f495413abc9cec3499e5eca7), [MSV000093168](https://massive.ucsd.edu/ProteoSAFe/dataset.jsp?task=c57d9dfc106c47b4983a093a63afa942). Other untargeted metabolomics dataset reanalyzed in the context of this study, which data was previously generated, include: [MSV000084062](https://massive.ucsd.edu/ProteoSAFe/dataset.jsp?task=7a35d515ea3f42039bddeb75b8d4ac1f), [MSV000096884](https://massive.ucsd.edu/ProteoSAFe/dataset.jsp?task=7a35d515ea3f42039bddeb75b8d4ac1f), [MSV000088645](https://massive.ucsd.edu/ProteoSAFe/dataset.jsp?task=6d079e4fb6874ebcadae21bf097ea2ed), and [MSV000095383](https://massive.ucsd.edu/ProteoSAFe/dataset.jsp?task=672ce0cd8cf3460cbfaa187f916cb0b9). Metagenomics sequencing data are available through EBI/ENA via the following accession codes: ERP165389 (3xTg) and ERP165388 (5xFAD). Additional information and processing pipelines are available via Qiita under the following study IDs: [3xTg Study ID #14748](https://qiita.ucsd.edu/) and [5xFAD Study ID #14957](https://qiita.ucsd.edu/study/description/14957).
- Code and data used for the untargeted metabolomics analysis are available on GitHub at https://github.com/simonezuffa/Manuscript_AD_Tissue. Code for tissuMASST and its associated web app are available at https://github.com/robinschmid/microbe_masst and https://github.com/mwang87/GNPS_MASST respectively. Code for the microbiome and multi-omic analyses is also available on GitHub and can be found at https://github.com/knightlab-analyses/U19_3xtg_5xfad_mouse_models.

## ACKNOWLEDGMENTS

Metabolomics data were generated by the Alzheimer’s Gut Microbiome Project (AGMP) or the Alzheimer’s Disease Metabolomics Consortium (ADMC) funded wholly or in part by the following grants and supplements thereto: NIA U19AG063744, U01AG061359, and R01AG081322 awarded to Dr. Kaddurah-Daouk at Duke University in partnership with a large number of academic institutions. As such, the investigators within the AGMP and the ADMC, not listed specifically in this publication’s author’s list, provided data along with its pre-processing and prepared it for analysis, but did not participate in analysis or writing of this manuscript. A listing of AGMP Investigators can be found at https://alzheimergut.org/meet-the-team/. A complete listing of ADMC investigators can be found at: https://sites.duke.edu/adnimetab/team/. The authors would like to thank technicians Caitlin Tribelhorn, Martin Casas Maya, Sawyer Farmer, and Gail Ackermann for their assistance with metagenomic sample processing.

## AUTHORCONTRIBUTIONS

S.Z. conducted untargeted metabolomics analysis and wrote the manuscript.

C.A. conducted metagenomics analysis and wrote the manuscript.

V.C.-L. and A.M.C.-R. run untargeted metabolomics experiments.

A.P., I.M., J.A., D. S. synthesized, purified, and performed MS/MS and RT matching of standards.

J.W.B., T.J.C., T.T, B.N., and M.deC.F. conducted animal experiments and collected samples.

R.S.B., L.H., H.T., and J.C. processed the samples for metagenomics experiments.

K.K., T.B., S.M., L.S., A.K.-P., S.F.G., D.S. provided feedback on metabolomics analysis and human cohorts.

M.W. developed Fast Search and infrastructure for tissueMASST.

R.K., R.K.-D., P.C.D., and S.K.M. designed the study, provided feedback, and secured funding. All authors reviewed and approved the manuscript.

## STAR METHODS

### Animal Study

As previously described^55^, 8-weeks-old wild-type C57BL/6J mice (The Jackson Laboratory, Cat#000664), homozygous 3xTg^22^ [B6;129-Tg(APPSwe,tauP301L)1Lfa *Psen*1^tm1Mpm^/Mmjax] (The Jackson Laboratory, Cat#004807) and hemizygous 5xFAD^23^ [B6.Cg-Tg(APPSwFlLon,PSEN1*M146L*L286V)6799Vas/Mmjax] (The Jackson Laboratory, Cat#034848) mice were obtained from the Jackson Laboratory and maintained in the laboratory of S.K.M at the California Institute of Technology (Caltech). Mice for 3xTg experiments were produced by homozygous breeding of the transgenic strain (Cat#004807) and homozygous breeding of wildtype (C57BL/6J, Cat#000664) control animals. 5xFAD hemizygous mice and wild-type littermates were produced by crossing transgenic mice with C57BL/6J mice. C57BL/6J, 3xTg, and 5xFAD mice were rederived as germ-free (GF) in the Caltech gnotobiotic facility. After weaning, mice were housed together by genotype with littermates and aged until the experiment dates. All experiments were performed with female and male mice. Mice underwent behavioral experiments, which are reported in Bostick et al.^55^ and GF mice were treated with antibiotics during behavioral tests [ampicillin (1 mg/mL), vancomycin (0.5 mg/mL), and neomycin (1 mg/mL)] to maintain the GF condition. Experimental mice were housed in sterilized microisolator cages and maintained on *ad libitum* autoclaved 5010 PicoLab Rodent Diet (LabDiet, Cat#5010) and sterilized water. Animals were group housed (2–5 mice per cage) unless otherwise specified. Ambient temperature in the animal housing facilities was maintained at 21-24°C, 30-70% humidity, with 13 h light / 11 h dark cycle. Detailed information on immune cell isolation and counting can be found in the companion paper by Bostick et al.^55^. All experiments were performed with approval from the Institutional Animal Care and Use Committee (IACUC) of Caltech (Protocol IA20-1798).

### Sample Collection

Fecal pellets were collected from each cage monthly and stored in Eppendorf tubes at -80 °C until use. Brain, liver, serum, and cecal contents were sampled following sacrifice at 7, 12 and 15 months for the 3xTg study, and at 5 and 8 months for the 5xFAD study. Each study included approximately 7-10 animals per genotype, colonization condition, sex, and timepoint. At sacrifice, mice were euthanized with CO_2_. Trunk blood was collected by cardiac puncture with a 1 mL Luer Lock syringe (BD Cat# 30968) and 25 G needle (BD Cat# 305122), deposited in a serum separator tube (Sarstedt Cat# 41.1378.005), and allowed to clot at room temperature for 15 min. The serum was separated by centrifugation at 10,000 x g for 5 min, transferred to an Eppendorf tube, flash frozen on dry ice, and stored at -80 °C until use. Tissues (whole brain, whole liver, cecum) were dissected. Brains were split at the midline and each hemisphere placed in a separate 1.5 mL Eppendorf tube, flash frozen on dry ice, and stored at -80 °C until use. Livers were placed in 5 mL Eppendorf tubes, flash frozen, and stored at -80 °C. Before distribution to analysis sites (Duke University and University of California San Diego), liver samples were thawed at room temperature, homogenized with a mortar and pestle pre-cooled with liquid nitrogen, divided between fresh Eppendorf tubes, flash frozen on dry ice, and stored at -80 °C until use. Contents were collected from the cecum into an Eppendorf tube, flash frozen on dry ice, and stored at -80 °C. Before distribution to analysis sites, cecal content samples were thawed at room temperature, homogenized with sterilized toothpicks, divided between fresh Eppendorf tubes, flash frozen on dry ice, and stored at -80 °C until use. Samples from 8-month-old male 5xFAD animals (SPF and GF) were affected by a freezer failure that may have compromised their quality, and this cohort was therefore excluded from analysis.

### Untargeted Metabolomics Sample Preparation

All solvents used during extraction and analysis were LC-MS grade (H_2_O Fisher W6-4; MeOH Fisher A456-4). Before extraction, samples were weighed and thawed at room temperature for 30 min to 1 h and extracted using cold 50% MeOH/H_2_O at a ratio of 100 mg of sample to 800 µL of solvent. A 5 mm stainless steel bead was added to the sample and homogenized in a Qiagen TissueLyser II for 5 min at 25 Hz. Samples were then incubated at 4 °C for 30 min before centrifugation at 15,000 x g for 15 min. The supernatant (200 µL) was transferred to a 96-well plate and dried in a CentriVap overnight before being stored at -80 °C until LC-MS/MS analysis. Serum samples were extracted using the Phree kit (Catalog no. 8E-S133-TGB Phenomenex Phree Phospholipid Removal). Fifty microliters were loaded onto the Phree 96-well plate. MeOH (100%) was added to the sample at a 4:1 ratio (MeOH:sample) and gently homogenized. The samples were centrifuged at 500 x g for 5 min, dried in the CentriVap overnight, and stored at -80 °C until LC-MS/MS analysis. All sample types were resuspended in 200 µL of 50% MeOH/H_2_O with an internal standard and incubated at –20 °C overnight for protein precipitation. Samples were centrifuged at 2000 rpm for 10 min and 150 µL were transferred to a shallow 96-well plate. Five microliters were collected from each sample per tissue type to create pooled quality controls (QCs).

### Untargeted Metabolomics Data Acquisition

Samples were randomized within each tissue (brain, serum, liver, cecal content, GF feces, 3xTg SPF feces, and 5xFAD SPF feces) and run in batches via an untargeted metabolomics pipeline comprising a Vanquish UHPLC (ultra-high performance liquid chromatography) system coupled to a Q-Exactive Orbitrap mass spectrometer (Thermo Fisher Scientific). Chromatography was performed using a Phenomenex C18 column (1.7 µm particle size, 2.1 mm x 50 mm) and a mobile phase of solvent A (water + 0.1% formic acid) and solvent B (acetonitrile + 0.1% formic acid). Injections of 5 μL of sample, with a flow rate of 0.5 mL/min, used the following gradient: 0-1 min 5% B, 1-7 min 5-99% B, 7-8 min 99% B, 8-8.5 min 99-5% B, 8.5-10 min 5% B. Tandem mass spectrometry (MS/MS) data was acquired via data-dependent acquisition (DDA) mode using exclusively positive electrospray ionization (ESI+). Briefly, the ESI parameters were set as follows: 53 L/min sheath gas flow, 14 L/min aux gas flow rate, 3 L/min sweep gas flow, 3.5 kV spray voltage, 269 °C inlet capillary, and the aux gas heater set to 438 °C. The MS scan range was set to 100 – 1500 *m/z* with a resolution at *m/z* 200 set to 35,000 with 1 microscans. Automatic gain control (AGC) was set to 5e4 with a maximum injection time of 50 ms. Up to 5 MS/MS (TopN = 5) spectra per MS1 were collected with a resolution at m/z *200* set to 17,500 with 1 microscan. Injection time was 50 ms with an AGC target of 5e4. The isolation window was set to 2.0 *m/z*. Normalized collision energy was set to a stepwise increase of 20, 30, and 40 eV with an apex trigger set to 2-15 s and a dynamic exclusion of 10 s.

### Untargeted Metabolomics Data Processing

Generated .raw files were converted into .mzML open-access format using ProteoWizard MSConvert^113^ and deposited in [GNPS/MassIVE](https://massive.ucsd.edu/ProteoSAFe/static/massive.jsp) under the following accession codes: [MSV000096035](https://massive.ucsd.edu/ProteoSAFe/dataset.jsp?task=b467af929b9347b8b0aa13f41957fa4d), [MSV000096036](https://massive.ucsd.edu/ProteoSAFe/dataset.jsp?task=8249c40f7e3d4ea8a4e20871725a7ca3), [MSV000096037](https://massive.ucsd.edu/ProteoSAFe/dataset.jsp?task=60905386a3f944e1b859b8d96592980b), [MSV000096038](https://massive.ucsd.edu/ProteoSAFe/dataset.jsp?task=7d99b115a6d346d5b787c7ce10030a79), [MSV000096039](https://massive.ucsd.edu/ProteoSAFe/dataset.jsp?task=55ec7f9e84124ca0827f0d7958192c34), [MSV000093230](https://massive.ucsd.edu/ProteoSAFe/dataset.jsp?task=fbf55f75f495413abc9cec3499e5eca7), [MSV000093168](https://massive.ucsd.edu/ProteoSAFe/dataset.jsp?task=c57d9dfc106c47b4983a093a63afa942). Feature detection and extraction were performed within each dataset using the batch processing mode in MZmine 3.9^114^. The .xml files used for batch processing can be found in the associated GitHub page. Briefly, data was imported using MS1 and MS2 detector via factor of lowest signal with noise factors set to 3 and 1.1, respectively. Sequentially, mass detection was performed and only ions acquired between 0.5 and 8 min, with MS1 and MS2 noise levels set to 5e4 and 1e3, respectively. Chromatogram builder parameters were set at 5 minimum consecutive scans, 1e5 minimum absolute height, and 10 ppm for *m/z* tolerance. Smoothing was applied before local minimum resolver, which had the following parameters: chromatographic threshold 85%, minimum search range retention time 0.2 min, minimum ratio of peak top/edge 1.7. Then, 13C isotope filter and isotope finder were applied. Features were aligned using join aligner with weight for *m/z* set to 3 and retention time tolerance set to 0.2 min. Features not detected in at least 3 samples were removed before performing peak finder. Ion identity networking and metaCorrelate were performed before exporting the final feature table. The GNPS^26^ and SIRIUS^53^ export functions were used to generate the feature table containing the peak areas and the .mgf files (MS/MS spectra list in text format) necessary for downstream analyses. Generated mzmine output per tissue, comprising a feature table and two separate .mgf files, one for molecular networking in GNPS and one for MS/MS molecular class prediction in SIRIUS are available on GitHub. Molecular classes of features with parent ion < 800 *m/z* were predicted via CANOPUS^54^ using SIRIUS 5.8^53^.

### Molecular Networking

Feature based molecular networking (FBMN)^115^ was performed in GNPS2^26^ for each tissue type. Networking parameters were set as follows: 0.02 for both precursor and fragment ion tolerances; 0.7 minimum cosine score and 5 minimum matching peaks; 10 Top K and 100 Max Component Size; and library search was set at 0.7 minimum cosine score and 4 minimum matching peaks. Library search was performed against GNPS libraries and the bile acid propagated library^116^. Data are available for download at the following links:

- Brain: https://gnps2.org/status?task=07339474d63641fe87bf8c371a1df6e8
- Serum: https://gnps2.org/status?task=2f80bbd389404bbfa4167504f8828593
- Liver: https://gnps2.org/status?task=b4b4d126c7524e238fa33561dd7fcf64
- Cecal content: https://gnps2.org/status?task=30f29bb9baf14c34bda7df559186af92
- Feces GF animals: https://gnps2.org/status?task=a2a90b1d952e4c41acbb11494632a7a3
- Feces SPF 3xTg study: https://gnps2.org/status?task=410d89ffe6a34003b234c6c21b2e5180
- Feces SPF 5xFAD study: https://gnps2.org/status?task=c806fe06c739445bbd708f8f6b9dfb65

Additionally, a cross-tissue molecular network was generated from the obtained tissue-specific .mgf files by a classical molecular network with the clustering function turned off. Networking parameters were the same as those used for FBMN and Min Cluster Size was set to 0. Output used for feature tracking was clusterinfo.tsv, clustersummary_with_network.tsv, and merged_pairs.tsv. Mapping of identical features across tissues was done by filtering the merged_pairs.tsv based on the following criteria: difference in parent mass < 0.02, difference in retention time (RT) < 0.2 min, modified cosine score > 0.9, and removing matches within the same tissue. The interactive cross-tissue molecular network can be downloaded at the following link:

- Cross-tissue network: https://gnps2.org/status?task=6c00d991f3954e3c984e3a3de9353cea

### tissueMASST

Building on the previously published tools microbeMASST^60^ and plantMASST^61^, tissueMASST was developed to capture all publicly available LC-MS/MS data acquired from human and rodent tissue samples deposited not only in GNPS/MassIVE, but also in MetaboLights^62^ and Metabolomics Workbench^63^; the two other major public repositories for metabolomics data. Through the efforts in controlled metadata reunification, 687,585 LC-MS/MS files were listed in [PanReDU](https://redu.gnps2.org/)^69^ in January 2025. Filtering the PanReDU table for datasets acquired from *Mus musculus*, *Rattus norvegicus*, and *Homo sapiens* via LC-MS/MS returned 153,508 files, of which 44,992 had accessible LC-MS/MS data. Of these, 27,954 files were human samples, 11,422 mouse samples, and 1,328 rat samples. These files were then used to generate the reference database of tissueMASST. Associated metadata information was also extracted and used to generate an interactive tree. Specifically, tissue type information (UBERONBodyPartName), species (NCBITaxonomy), health status (DOIDCommonName), sex (BiologicalSex), and age (LifeStage) were used. Dash and Flask open-source libraries were used to build a tissueMASST web application available at https://masst.gnps2.org/tissuemasst/ and associated code available on GitHub (https://github.com/mwang87/GNPS_MASST/blob/master/dash_tissuemasst.py). Researchers can use the web app to search for MS/MS spectra via a Universal Spectrum Identifier (USI)^117^ or by manually inputting the fragment ions and their intensities. Batch searches are enabled via a custom Python script (https://github.com/robinschmid/microbe_masst), which can use as input a tabular file (.tsv or .csv) containing a list of USIs or a single .mgf file containing the MS/MS of interest. As previously described^60^, tissueMASST also leverages the Fast Search API (https://fasst.gnps2.org/fastsearch/) to search MS/MS spectra of interest against all the MS/MS spectra that have been indexed across GNPS/MassIVE, MetaboLights and Metabolomics Workbench. Search outputs are visualized as interactive trees that can be downloaded as HTML files and summary tables.

### Untargeted Metabolomics Data Analysis

Tissue-specific feature tables were imported into R 4.2.2 (R Foundation for Statistical Computing, Vienna, Austria) and analyzed separately. Data quality was assessed by inspecting total extracted peak areas, and coefficients of variance (CV) of internal standard (IS) in the biological samples, and 6 different standards (Amitriptyline, Sulfadimethoxine, Sulfamethazine, Sulfamethizole, Sulfachloropyridazine, and Coumarin 314) present in the QCmix samples, which were run every 10 biological samples. Blank subtraction was performed by removing features detected in Blank and QCmix samples whose mean peak areas were not at least 5 times the ones observed in the QCpool. Feature tables were robust center log ratio (RCLR) transformed using ‘vegan v 2.6’ and features with near-zero variance were removed using ‘caret v 6.0’ before performing dimensionality reduction. Principal component analysis (PCA) and partial least squares discriminant analysis (PLS-DA) were performed using mixOmics ‘v 6.22’. PERMANOVA was used to evaluate group centroid separation in PCA using ‘vegan v 2.6’. PLS-DA model performances were obtained via 5-folds cross-validation and 999 permutations. Features important for group separation were identified via variable importance (VIP) scores > 1. Peak areas of features associated with different conditions (*e.g*., AD vs WT) obtained from PLS-DA models were collapsed to generate ratios which were then log transformed before testing for significant differences using Wilcoxon Test. Linear mixed-effect models were obtained via ‘lmerTest v 3.1’ and animal id was used as a random effect. Detection incidence of the unknown carnitines in human data was tested via Chi-squared test. UpSet plots were generated using ‘UpSetR v 1.4’. The packages ‘tidyverse v 2.0’ and ‘ggpubr v 0.6’ were used for data manipulation and visualization. Code for the analysis and to generate the figures is available on GitHub (https://github.com/simonezuffa/Manuscript_AD_Tissue).

### Re-Analysis of External Untargeted Metabolomics Datasets

Four external untargeted metabolomics datasets were investigated to extract and analyze the carnitines of interest. Analyzed data were collected either from serum or plasma. One dataset ([MSV000084062](https://massive.ucsd.edu/ProteoSAFe/dataset.jsp?task=abbb4996c26843f9a8c73eba0d743794)) was acquired from C57BL/6J mice exposed to either a cocktail of antibiotics, inulin, or standard chow diet. The other three datasets ([MSV000096884](https://massive.ucsd.edu/ProteoSAFe/dataset.jsp?task=7a35d515ea3f42039bddeb75b8d4ac1f), [MSV000088645](https://massive.ucsd.edu/ProteoSAFe/dataset.jsp?task=6d079e4fb6874ebcadae21bf097ea2ed), and [MSV000095383](https://massive.ucsd.edu/ProteoSAFe/dataset.jsp?task=672ce0cd8cf3460cbfaa187f916cb0b9)) were acquired from humans and are part of the [Alzheimer Gut Microbiome Project](https://alzheimergut.org/) (AGMP) Consortium and [Accelerating Medicines Partnership for Alzheimer Disease](https://fnih.org/our-programs/accelerating-medicines-partnership-amp/amp-alzheimers-disease-2-0/) (AMP-AD). Open-source format files .mzML and .mzXML were processed via MZmine 3.9 as previously described using the same parameters for all datasets. From the generated quantification tables, per sample TICs were calculated. Outliers with TIC below or above the first or third quartile ± 1.5 times the interquartile range (IQR) were excluded from downstream analysis. Peak areas of the carnitine of interest were extracted and log transformed. Metadata for the human datasets, ADRC, T-1000, and MARS, is available via the [AD Knowledge Portal](https://adknowledgeportal.synapse.org/) upon request. Briefly, the ADRC cohort includes AGMP participants recruited from multiple NIA-funded ADRCs across the U.S., who consented to participate in the observational longitudinal study at their respective ADRCs. Collected neuropsychological data, data on depression, anxiety, medications, medical history, and additional metadata followed NACC Uniform Data Set (UDS) 3. Participants donated blood during annual visits to the clinic and were provided a home collection kit in person or by mail to collect a small amount of fecal material. If multiple blood samples were collected from an individual, only the most recently acquired sample was included in this analysis. In total, 490 distinct samples were included in the analysis. The ADRC population characteristics were as follows: 61.59% female; 78.66% white; mean age 72.28±7.87 (SD) years; mean BMI 27.12±5.29 (SD); 12.20% affected by type 2 diabetes; 6.70% affected by depression; 25.81% diagnosed with MCI or dementia; and 18.69% with a presumptive diagnosis of AD. Clinical diagnosis of mild cognitive impairment (MCI) or dementia and presumptive diagnosis of AD were determined by a multidisciplinary consensus diagnostic panel based on the National Institute on Aging–Alzheimer’s Association (NIA-AA) criteria^118,119^. Memory performance was assessed using the Craft Story 21 Recall (Delayed) test, scored by the total number of story units recalled verbatim. Descriptions of sample collection and population characteristics for the T-1000 and MARS cohorts are available in the respective publications by Dilmore et al.^66^ and Kang et al.^67^. Similarly for these cohorts, if repeated measures from the same individual were available, only the latest sample was considered. Constructed models were adjusted for age, sex, and BMI unless otherwise specified. Feature based molecular networks with associated feature tables and annotations are available for download at the following links:

- Animal: https://gnps2.org/status?task=dca12e98916e4b699eb625c92faff33f
- ADRC: https://gnps2.org/status?task=c9cdd788241e468fb7c6a1fecc1a7bce
- T-1000: https://gnps2.org/status?task=1539c7aa8feb4c2da695009900bac943
- MARS: https://gnps2.org/status?task=e9ce7b4d883b4c2eae7b0ca8be48ace2

### Synthesis of Carnitine Standards

L-Carnitine hydrochloride (CAS Number: 6645-46-1), phenylpropionyl chloride (CAS Number: 645-45-4), phenylacetyl chloride (CAS Number: 103-80-0), pyridine (CAS Number: 110-86-1) and benzoyl chloride (CAS Number: 98-88-4) were purchased from Sigma Aldrich. L-Carnitine hydrochloride (2.53 mmol, 500 mg, 1 eq.) was dissolved in 4 mL dry dichloromethane (DCM) in a 20 mL scintillation vial with a magnetic stir bar and placed in an ice bath at 0 °C. To this solution, (5.06 mmol, 409 µL, 2 eq.) dry pyridine was added, and the equivalent amount of acyl chloride was slowly added at 0 °C while stirring. The reaction was allowed to stir for 1–3 h at room temperature, with progress monitored for completion. After the reaction was complete, the solvent was removed by rotary evaporation, and the synthesized compounds were directly used for further analysis. Retention time and MS/MS matching of the generated standards (benzoyl-carnitine, phenylacetyl-carnitine, and phenylpropionyl-carnitine) to the animal samples (3xTg) and human samples (ADRC, T-1000, and MARS) was performed via UHPLC-MS/MS analysis using the same settings as described for the acquisition of the 3xTg and 5xFAD samples. Standards were also spiked in for Level 1 annotation.

### Shotgun Metagenomics Data Acquisition and Processing

Fecal samples (0.03-0.04 g) were transferred into 1 mL Matrix Tubes (ThermoFisher, Waltham, MA, USA) before gDNA extraction. Samples were then extracted for microbiome sequencing using reagents from the MagMAX Microbiome Ultra Nucleic Acid Isolation Kit (ThermoFisher Scientific, Waltham, MA, USA). The protocol was adapted in order to perform the lysis and bead beating extraction steps in the 1 mL Matrix Tube sample collection devices, eliminating the sample-to-bead plate transfer step and minimizing potential well-to-well cross-contamination during bead beating^27^. After bead beating, samples were transferred to KingFisher plates (ThermoFisher, Waltham, MA, USA) and gDNA was extracted according to kit instructions. Extracted gDNA was quantified using the Quant-iT PicoGreen dsDNA Assay Kit (Invitrogen, Waltham, MA, USA) and normalized to 5 ng in 3.5 μL sterile water for library preparation, which was performed using a miniaturized adaptation of the KAPA HyperPlus Library Kit (Roche, Basel, Switzerland)^120^. The library was quantified via PicoGreen Assay (Invitrogen), and all samples were equal volume pooled, PCR cleaned (QIAGEN), and size selected from 300-700 bp using a Pippin HT (Sage Sciences). QC was run on an Agilent 4000 Tapestation (Agilent, Santa Clara, CA, USA) to confirm expected library sizes after PCR cleanup and size selection. The equal volume pool was sequenced on an iSeq100 (Illumina, San Diego, CA, USA). Utilizing the sample concentration and read counts per sample obtained from the iSeq100 run, a normalized pooling value was calculated for each sample to optimize multiplexing efficiency and achieve a more even read count distribution during NovaSeq sequencing^121^. After re-pooling the library with the iSeq normalized pool values, samples were PCR cleaned, size selected (300-700 bp), and QC was performed using an Agilent tapestation. The PCR-cleaned, size-selected, iSeq-normalized pool was then sequenced on a NovaSeq 6000 (Illumina, San Diego, CA, USA) at the Institute for Genomic Medicine at the University of California, San Diego with a S4 flow cell and 2×150bp chemistry.

### Shotgun Metagenomics Data Analysis

Raw sequence reads (BCL files) were demultiplexed to per sample FASTQ and quality filtered^122^. Adapter trimming was performed by fastp^123^ and the resulting FASTQ files were uploaded into Qiita^124^. Metagenomic reads were host filtered (*Mus musculus*) using the C57BL/6J reference genome GCF_000001635.27 (GRCm39). Host filtered files were then processed with the default shotgun metagenomic analysis pipeline performed using Woltka (v0.1.4)^125^. In brief, direct genome alignments were made against the “Web of Life” database (release 2)^126^, which contains 10,575 microbial genomes of bacteria and archaea. The sequence alignment was performed using bowtie2^127^ and by mapping sequencing data to microbial reference genomes. Reads mapped to a microbial reference genome were counted as hits such that the resultant feature table comprises samples (rows) by microbial genome IDs (columns) and concomitant abundances. If a sequence mapped to multiple genomes by Bowtie2 (up to 16), each genome was counted 1/k times, where k is the number of genomes to which a sequence was mapped. Microbial genome IDs are considered operational genomic units (OGUs) and provide a shotgun metagenomic equivalent to amplicon sequence variants in 16S rRNA amplicon sequencing data^125^. The OGU frequencies were then summed after the entire alignment was processed and rounded to the nearest even integer, thereby making the sum of OGU frequencies per sample nearly equal (considering rounding) to the number of aligned sequences in the dataset. The resultant count matrix was saved as a .biom format table^128^. Functional profiles created by matching against Prodigal-estimated ORFs to gene ontology (GO) then cascading annotations utilizing Uniref and KEGG. The per-genome OGU biom file was downloaded and converted to a QIIME2^129^(v2024.5) artifact. For the 3xTg study, due to overpooling of blanks after the iSeq normalization step, samples with either low starting DNA concentration (including blanks) or poor amplification with less than 2,000 counts/pooled volume were dropped from further analysis. This was not necessary for the 5xFAD study. The feature table was rarified to a depth of 1,000,000 reads/sample for both the 3xTg and 5XFAD Studies to control for sequencing effort before performing alpha diversity analysis via Shannon Index^130^. For the log-based diversity metrics phylo-RPCA^25^, an unrarified table was used for calculations and downstream analyses. Greengenes2 (v2022.10) was used for taxonomy and phylogeny assignments^131^. ANCOM-BC2^132^ was used to determine species and functional pathway differential abundance^132^. Before performing differential abundance analysis, OGUs were collapsed at species level and only species present in at least 5% of the samples and accounting for at least 0.00001% of the total generated reads of the study were retained.

### Gene Search

Gene searches were specifically performed for *fldC*, a component of the phenyllactate dehydratase complex FldABC, and *bsh*, a bile salt hydrolase/transferase. For *fldC*, a multispecies version of the gene was used (https://www.ncbi.nlm.nih.gov/protein/WP_003492354.1/). For *bsh*, files were acquired from NCBI and Uniprot, pooled, and then amino acid sequences with at least 70% similar identity were clustered via USEARCH^133^ into a unique database. Then, DIAMOND^57^ blastx was used for read alignment and matching with parameter max-target-seqs 1 and a minimum 45% amino acid sequence identity. Metagenomic reads from each sample were aligned and matched to the genes of interest against the respective usearch clustered databases using Diamond blastx. Output dataframes were further filtered for e-value < 0.00001, > 20 bp of alignment length, and > 60% query alignment ratio before plotting.

### Multi-Omics Data Analysis

Joint-RPCA is an unsupervised method used to agnostically determine co-occurrences between the microbiome and metabolome data^25,58^. In brief, the microbiome and metabolome raw feature tables were first RCLR transformed. This helps to account for the sparsity and compositionality of microbiome and metabolomics data. The raw observed values were only computed on the non-zero entries and then averaged and optimized. The shared samples of all input matrices were used to estimate a shared matrix. The estimated shared matrix and the matrix of shared eigenvalues across all input matrices were recalculated at each iteration to ensure consistency. Minimization was performed across iterations by gradient descent. Cross-validation of the reconstruction was performed in order to prevent overfitting of the joint factorization. Co-occurrences of features across the two input matrices were calculated from the final estimated matrices.

## SUPPLEMENTAL INFORMATION

### Supplementary Figures. Figures S1–S9

**Figure S1.** Breakdown of Collected Samples

**Figure S2.** Acquired Untargeted Metabolomics Data per Tissue

**Figure S3.** Modeling Workflow for Metabolic Feature Categorization

**Figure S4.** Sex Influence on Log Ratios in the 3xTg Study

**Figure S5.** Sex Influence on Log Ratios in the 5xFAD Study

**Figure S6.** Additional Molecular Features Dysregulated in 3xTg Mice

**Figure S7.** Metagenomics Analysis of Other Covariates

**Figure S8.** Available Metadata for tissueMASST

**Figure S9.** Retention Time and MS/MS Matching of Synthetized Carnitine Standards

### Supplementary Tables. Tables S1-S22

**Table S1.** PLS-DA Discriminant Features in 3xTg SPF Brains

**Table S2.** PLS-DA Discriminant Features in 3xTg SPF Serum

**Table S3.** PLS-DA Discriminant Features in 3xTg SPF Liver

**Table S4.** PLS-DA Discriminant Features in 3xTg SPF Cecum

**Table S5.** PLS-DA Discriminant Features in 3xTg SPF Feces

**Table S6.** PLS-DA Discriminant Features in 3xTg GF Feces

**Table S7.** PLS-DA Discriminant Features in 5xFAD SPF Brains

**Table S8.** PLS-DA Discriminant Features in 5xFAD SPF Serum

**Table S9.** PLS-DA Discriminant Features in 5xFAD SPF Liver

**Table S10.** PLS-DA Discriminant Features in 5xFAD SPF Cecum

**Table S11.** PLS-DA Discriminant Features in 5xFAD SPF Feces

**Table S12.** PLS-DA Discriminant Features in 5xFAD GF Feces

**Table S13.** Joint-RPCA of Brain Metabolome and Fecal Microbiome in 3xTg SPF

**Table S14.** Joint-RPCA of Serum Metabolome and Fecal Microbiome in 3xTg SPF

**Table S15.** Joint-RPCA of Liver Metabolome and Fecal Microbiome in 3xTg SPF

**Table S16.** Joint-RPCA of Cecal Metabolome and Fecal Microbiome in 3xTg SPF

**Table S17.** Joint-RPCA of Fecal Metabolome and Fecal Microbiome in 3xTg SPF

