## Supplementary material for "A Multi-Organ Murine Metabolomics Atlas Reveals Molecular Dysregulations in Alzheimer’s Disease": Figure S1

| Study | Strain | Colonization | Sex | Age (months) | Number of Animals |
|---|---|---|---|---|---|
| 3xTg | 3xTg | GF | female | 7 | 6 |
| 3xTg | 3xTg | GF | male | 7 | 9 |
| 3xTg | 3xTg | SPF | female | 7 | 8 |
| 3xTg | 3xTg | SPF | male | 7 | 6 |
| 3xTg | WT | GF | female | 7 | 7 |
| 3xTg | WT | GF | male | 7 | 9 |
| 3xTg | WT | SPF | female | 7 | 7 |
| 3xTg | WT | SPF | male | 7 | 7 |
| 3xTg | 3xTg | GF | female | 12 | 9 |
| 3xTg | 3xTg | GF | male | 12 | 5 |
| 3xTg | 3xTg | SPF | female | 12 | 8 |
| 3xTg | 3xTg | SPF | male | 12 | 8 |
| 3xTg | WT | GF | female | 12 | 7 |
| 3xTg | WT | GF | male | 12 | 11 |
| 3xTg | WT | SPF | female | 12 | 8 |
| 3xTg | WT | SPF | male | 12 | 9 |
| 3xTg | 3xTg | GF | female | 15 | 11 |
| 3xTg | 3xTg | SPF | female | 15 | 7 |
| 3xTg | WT | GF | female | 15 | 9 |
| 3xTg | WT | SPF | female | 15 | 9 |
| 5xFAD | 5xFAD | GF | female | 5 | 14 |
| 5xFAD | 5xFAD | GF | male | 5 | 9 |
| 5xFAD | 5xFAD | SPF | female | 5 | 11 |
| 5xFAD | 5xFAD | SPF | male | 5 | 8 |
| 5xFAD | WT | GF | female | 5 | 11 |
| 5xFAD | WT | GF | male | 5 | 8 |
| 5xFAD | WT | SPF | female | 5 | 9 |
| 5xFAD | WT | SPF | male | 5 | 15 |
| 5xFAD | 5xFAD | GF | female | 8 | 8 |
| 5xFAD | 5xFAD | GF | male | 8 | 8 |
| 5xFAD | 5xFAD | SPF | female | 8 | 9 |
| 5xFAD | 5xFAD | SPF | male | 8 | 8 |
| 5xFAD | WT | GF | female | 8 | 5 |
| 5xFAD | WT | GF | male | 8 | 6 |
| 5xFAD | WT | SPF | female | 8 | 7 |
| 5xFAD | WT | SPF | male | 8 | 13 |

**Figure S1. Breakdown of Collected Samples**

On average, an equal number of male and female animals were investigated and sacrificed to collect tissues at each timepoint - 7, 12, and 15 months of age in the 3xTg study, 5 and 8 months of age in the 5xFAD study. For the 15 months timepoint in the 3xTg study, only female mice were investigated.

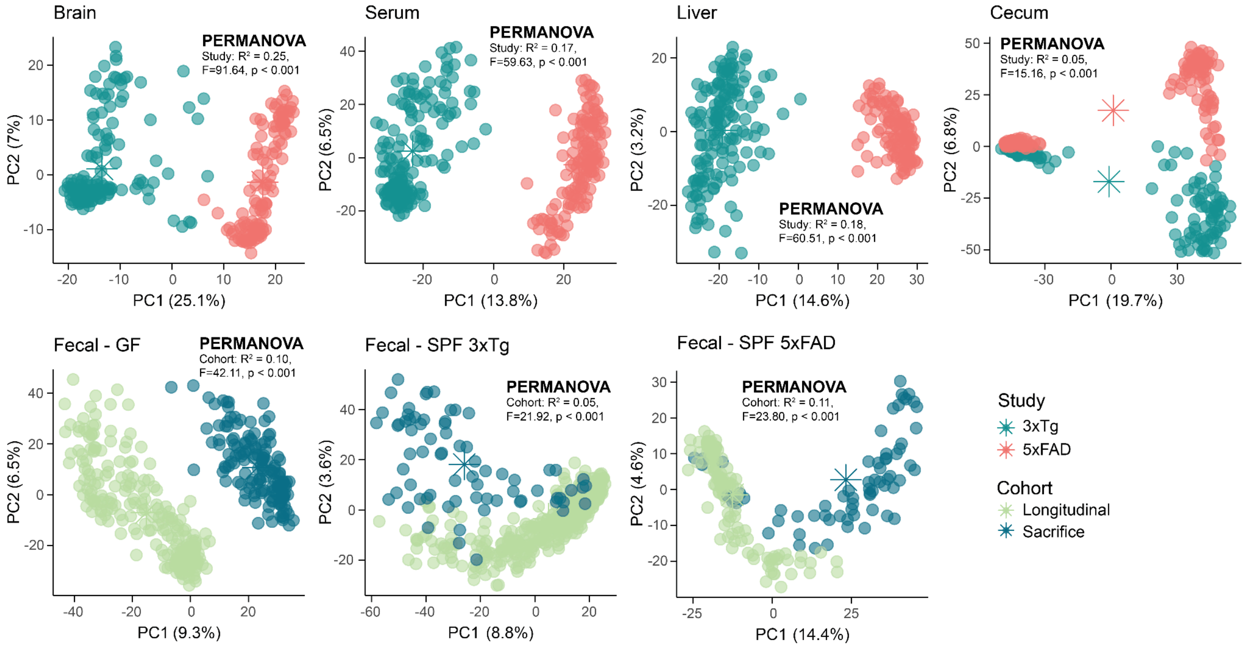


**Figure S2. Acquired Untargeted Metabolomics Data per Tissue**

Brain, serum, liver, and cecal samples metabolomics data from the 3xTg and 5xFAD studies were randomized and acquired on a single continuous run. Nevertheless, clear separation between metabolic profiles of samples belonging to the two different studies was observed (PERMANOVA, $p < 0.001$). Cecal content data included samples from both SPF and GF animals and colonization represented the greatest source of explained variance (PERMANOVA, $R^2 = 0.31$, $F = 131.9$, $p < 0.001$). Fecal sample data from the two different studies were acquired on three different runs (GF, SPF 3xTg, and SPF 5xFAD). In this case, a separation between samples belonging to Longitudinal or Sacrifice cohorts was observed (PERMANOVA, $p < 0.001$). Asterisks in the PCA score plots indicate group centroids.

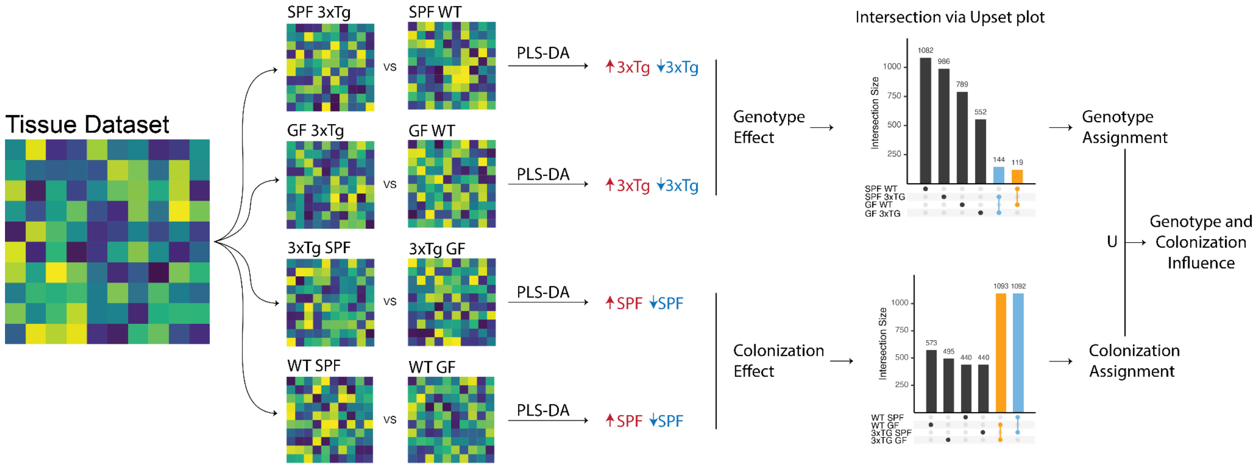


**Figure S3. Modeling Workflow for Metabolic Feature Categorization**

Each tissue dataset (brain, serum, liver, cecum, and feces) was stratified first by study (3xTg or 5xFAD) and then by genotype (3xTg/5xFAD or WT) or colonization (SPF or GF). Four PLS-DA models were then generated to extract information characterizing genotype (SPF_3xTg vs SPF_WT and GF_3xTg vs GF_WT) or colonization (3xTg_SPF vs 3xTg_GF and WT_SPF vs WT_GF) effect. From these models, metabolic features with variable importance score (VIP) > 1 were extracted and considered significant for group separation. These features were either increased or decreased in AD animals (3xTg or 5xFAD) compared to WT or increased or decreased in colonized animals (SPF) compared to GF. Upset plots were generated to identify shared metabolic features with same directionality within each study (*e.g.*, increased in SPF animals in both 3xTg and WT genotype background). This allows for a greater degree of confidence in the understanding if a metabolic feature was influenced or not by the genotype or the presence of a microbiome. Finally, categorized features were intersected to identify the ones that were affected by both conditions (genotype and colonization). The figure illustrates a representative schematic for cecal content data analysis in the 3xTg study. The same workflow was used for each tissue type and in the 5xFAD study. Red indicates features higher in 3xTg or SPF animals, while light blue indicates features lower in 3xTg or SPF.

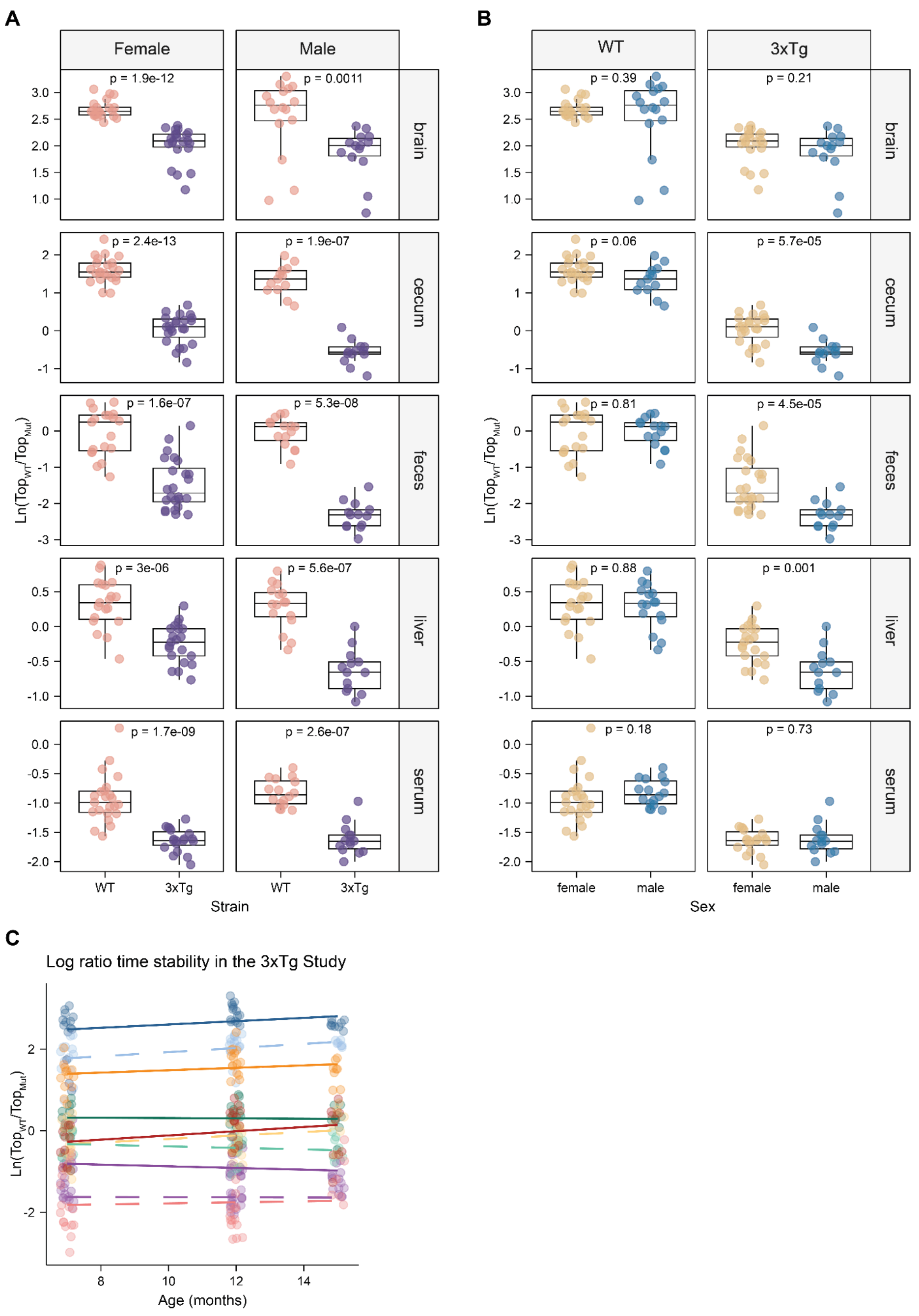


**Figure S4. Sex Influence on Log Ratios in the 3xTg Study**

**(A)** Sex stratification maintained significance in differential log ratios of metabolic features separating 3xTg from WT. **(B)** Within the 3xTg genotype only, a significant separation of the log ratios from the metabolic profiles of the liver, cecal content, and feces was observed between male and female. Ratio appeared to be exacerbated in male animals. Significance tested via Wilcoxon tests followed by BH correction. Boxplots show first (lower) quartile, median, and third (upper) quartiles.

**(C)** Linear regression on tissue log ratios through time did not reveal any significant influence of age in the 3xTg study. Dashed lines represent 3xTg animals while the solid ones WT (brain, blue; orange, cecal content; red, feces; green, liver; purple, serum).

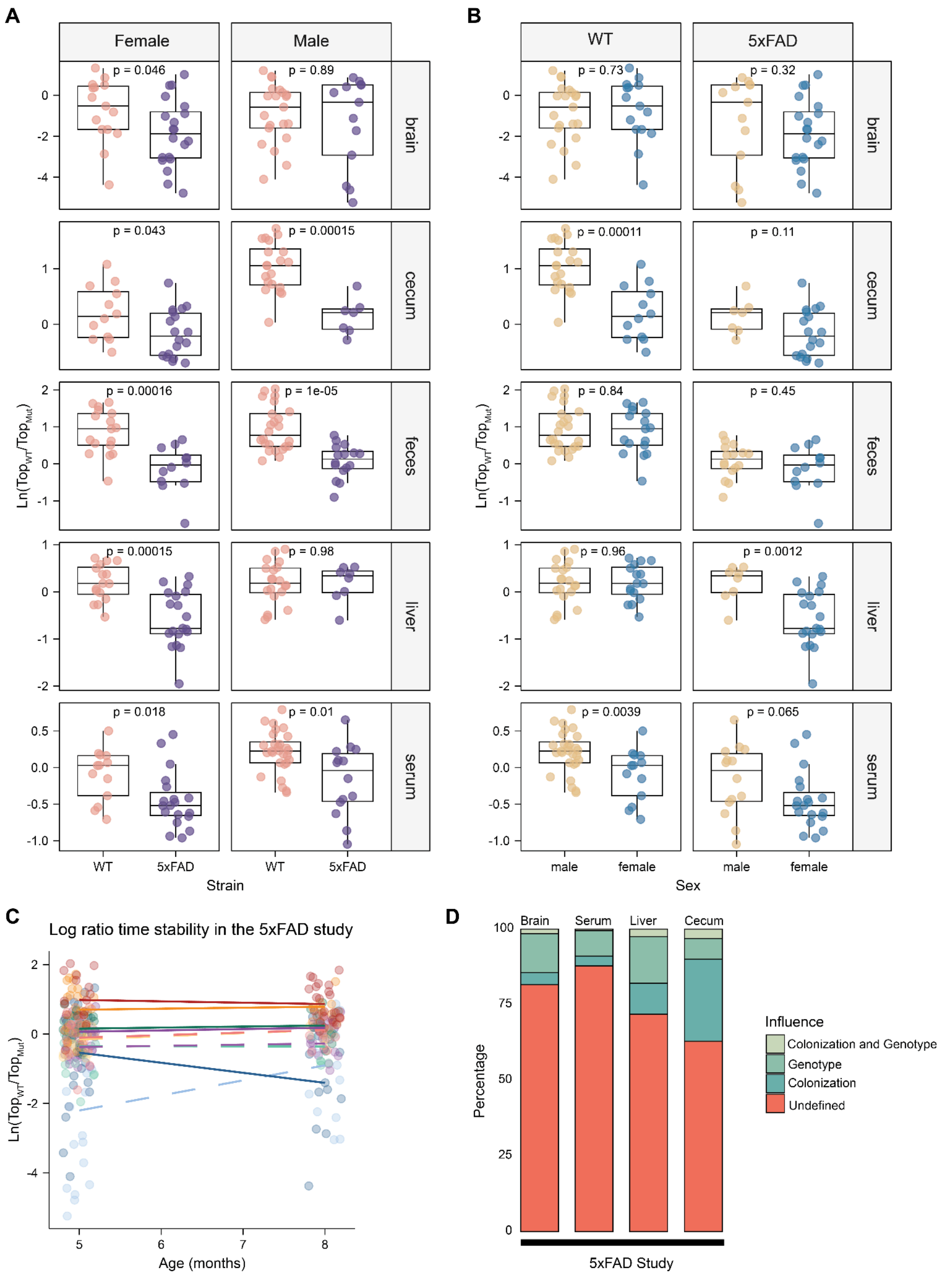


**Figure S5. Sex Influence on Log Ratios in the 5xFAD Study**

**(A)** Sex stratification disrupted significance of differential log ratios between 5xFAD and WT in male animals for liver and brain metabolic profiles.

**(B)** Sex differences of the log ratios were observed in both 5xFAD and WT for serum metabolic profiles, and in the cecum profile for WT and the liver for 5xFAD. Significance tested via Wilcoxon tests followed by BH correction. Boxplots show first (lower) quartile, median, and third (upper) quartiles.

**(C)** Ratios were not affected by age.

**(D)** Influence of colonization or genotype on discriminating features of interest extracted from PLS-DA models obtained via the nested workflow described in **Figure S3**. A stronger colonization effect was observed in cecum.

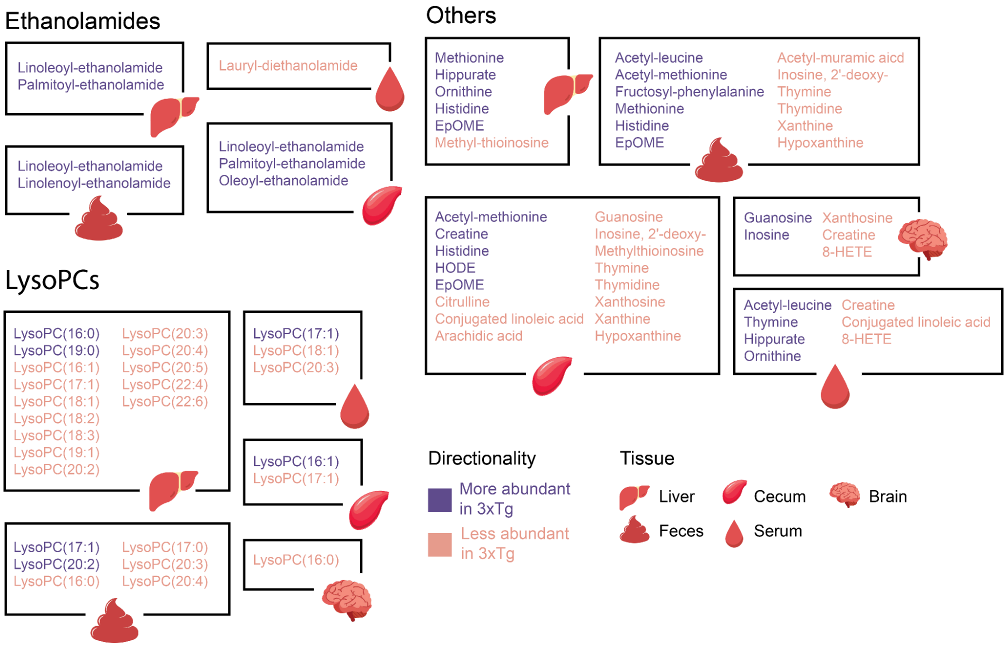


**Figure S6. Additional Molecular Features Dysregulated in 3xTg Mice**

Schematic representation of additional annotated molecules altered in 3xTg mice on a SPF background found via PLS-DA models. Ethanolamides, lysoPCs, and several other molecules including purine and pyrimidine derivatives, linoleic acid metabolites, metabolites part of the urea cycle, methionine, histidine, and hippurate were affected. Annotations are based on MS/MS spectral matching, which correspond to a Level 2 or 3 annotation according to the Metabolomics Standard Initiative (MSI).

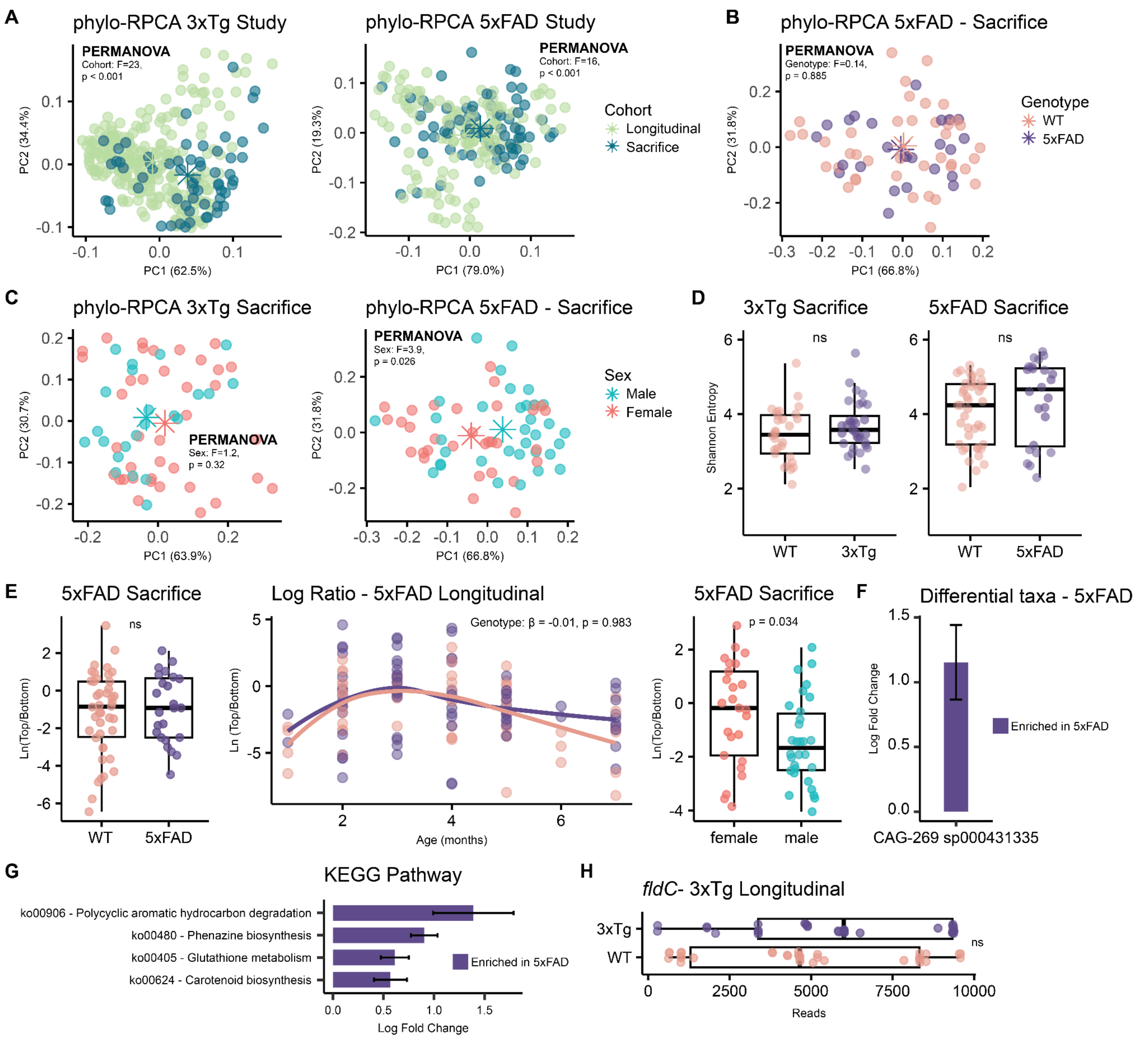


**Figure S7. Metagenomics Analysis of Other Covariates**

**(A)** Phylo-RPCA analysis of fecal samples from the 3xTg and 5xFAD studies reveals significant separation of samples from the Sacrifice and Longitudinal cohorts (PERMANOVA, $p < 0.001$) due to differences in sample collection.

**(B)** No differences based on genotype (5xFAD or WT) were observed in the 5xFAD Sacrifice samples

**(C)** Sex did not influence the fecal microbial profiles in the 3xTg study but it had a possible small effect in the 5xFAD.

**(D)** No difference in alpha diversity (Shannon Diversity Index) was observed based on genotype in both studies.

**(E)** No difference in possible log ratio of differential bacteria was observed in the 5xFAD Sacrifice samples. The defined log ratio was then applied to the Longitudinal data but also in this case no differences were observed. A significant log ratio was established to separate 5xFAD animals based on sex.

ANCOM-BC2 results for differential abundance taxa **(F)** and pathways **(G)** observed in the 5xFAD study. AD animals presented higher levels of CAG-269 and pathways involved in polycyclic aromatic hydrocarbon degradation, phenazine biosynthesis, and glutathione metabolism.

**(H)** No differences in encoded *fldC* gene levels between WT and 3xTg animals in the Longitudinal cohort. Importantly, *fldC* were not observed in the 3xTg Sacrifice cohort. Significance tested via Wilcoxon test.

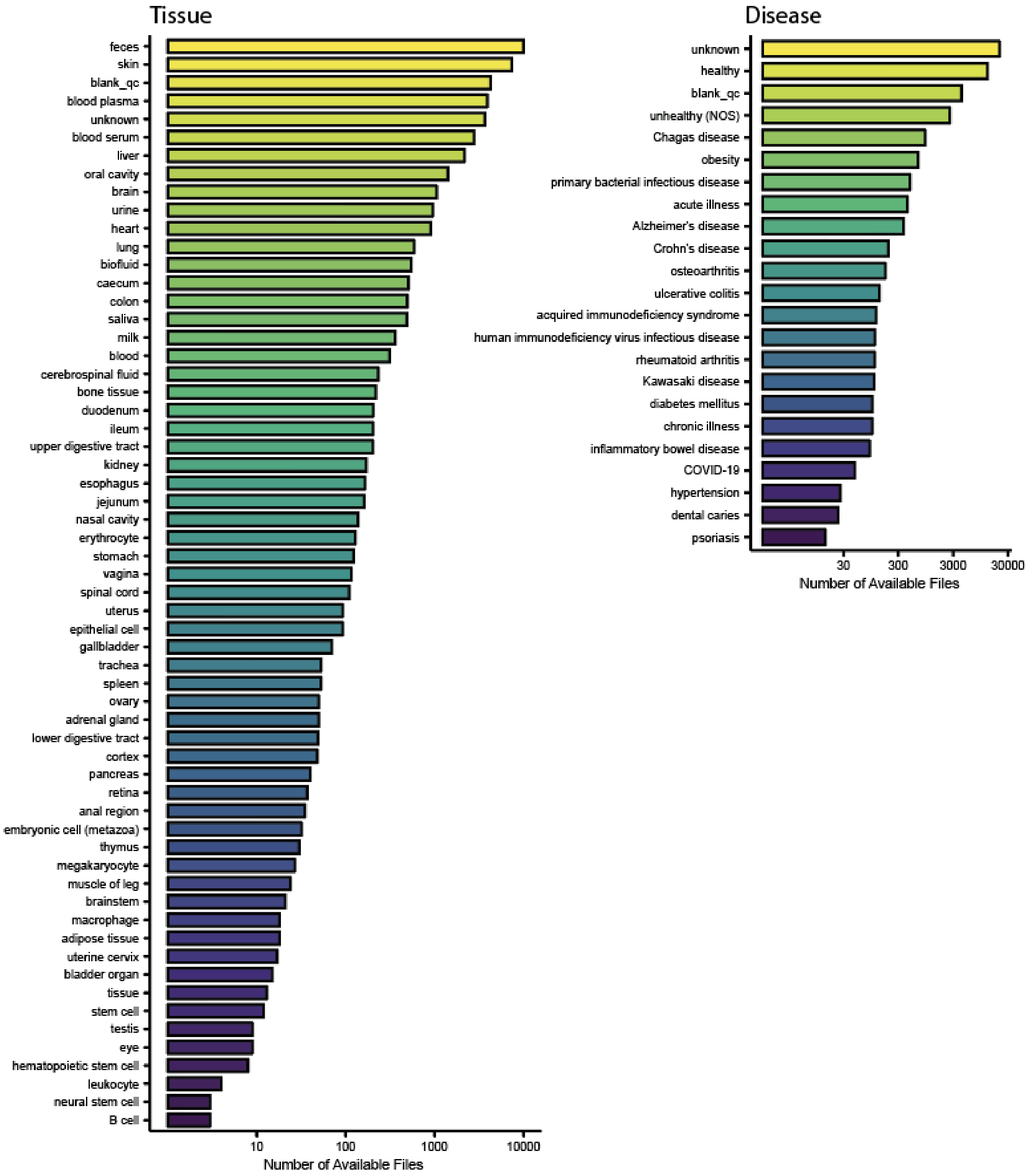


**Figure S8. Available Metadata for tissueMASST**

Number of files available for tissue and diseases metadata category across *M. musculus*, *R. norvegicus*, and *H. sapiens* samples present in the reference database of tissueMASST. Metadata information was acquired via PanReDU in January 2025. Availability of metadata depends on the specific dataset based on owner sharing policies. Public availability of MS data via MassIVE is not always reflected by metadata availability for the dataset of interest..

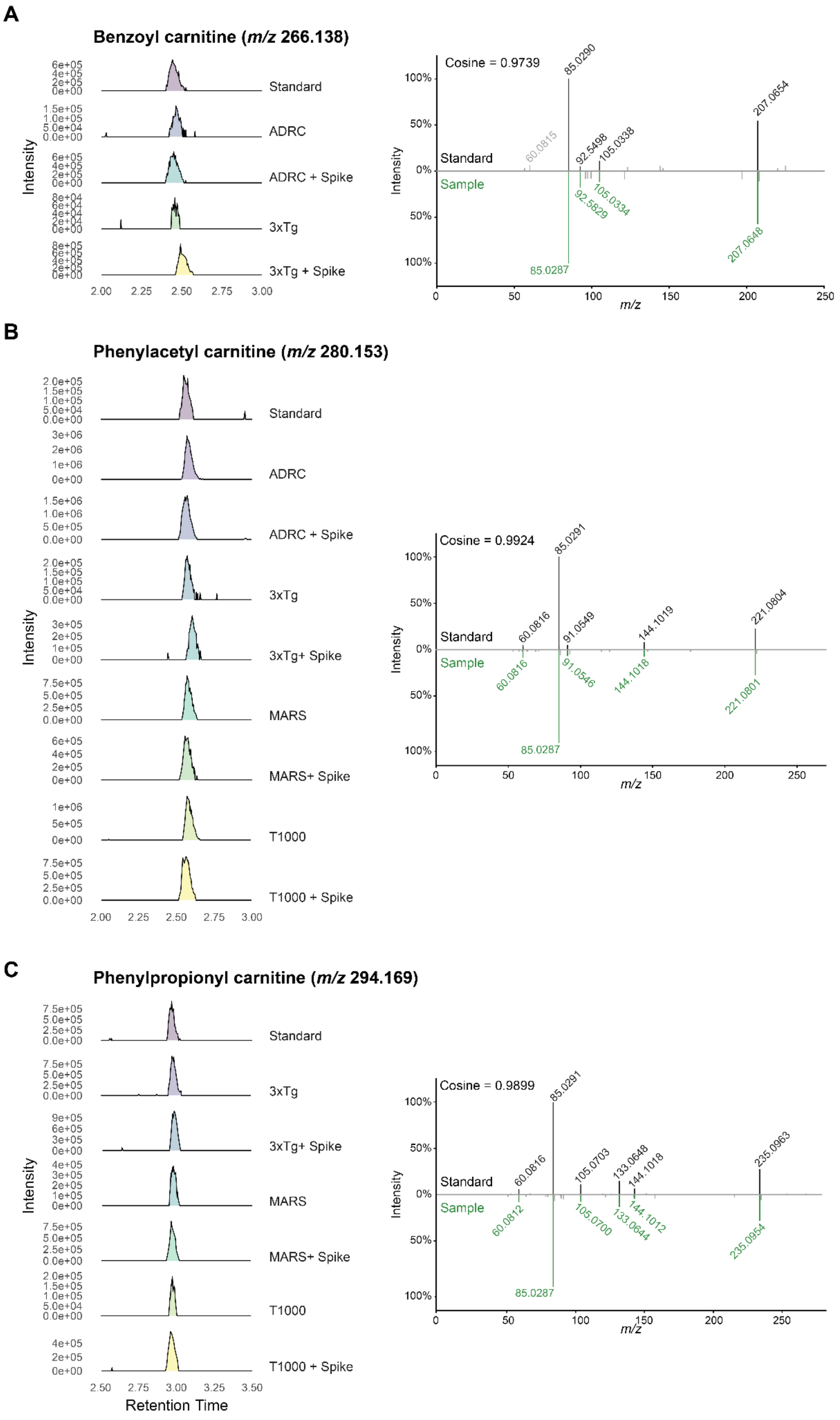


**Figure S9. Retention Time and MS/MS Matching of Synthetized Carnitine Standards**

Generated standards for benzoyl-carnitine (M+H, *m/z* 266.138), phenylacetyl-carnitine (M+H, *m/z* 280.153), and phenylpropionyl-carnitine (M+H, *m/z* 294.169) were acquired alongside the other animals (3xTg) and human (ADRC, T-1000, and MARS) samples and spiked-in to obtain a Level 1 annotation via MS/MS and retention time matching.
